# Stringent Response Governs the Virulence and Oxidative Stress Resistance of Francisella tularensis

**DOI:** 10.1101/653790

**Authors:** Zhuo Ma, Kayla King, Maha Alqahtani, Madeline Worden, Parthasarthy Muthuraman, Christopher Cioffi, Chandra Shekhar Bakshi, Meenakshi Malik

## Abstract

*Francisella tularensis* is a Gram-negative bacterium responsible for causing tularemia in the northern hemisphere. *F. tularensis* has long been developed as a biological weapon due to its ability to cause severe illness upon inhalation of as few as ten organisms and based on its potential to be used as a bioterror agent is now classified as a Tier 1 Category A select agent by the CDC. The stringent response facilitates bacterial survival under nutritionally challenging starvation conditions. The hallmark of stringent response is the accumulation of the effector molecules ppGpp and (p)ppGpp known as stress alarmones. The *relA* and *spoT* gene products generate alarmones in several Gram-negative bacterial pathogens. RelA is a ribosome-associated ppGpp synthetase that gets activated under amino acid starvation conditions whereas, SpoT is a bifunctional enzyme with both ppGpp synthetase and ppGpp hydrolase activities. *Francisella* encodes a monofunctional RelA and a bifunctional SpoT enzyme. Previous studies have demonstrated that stringent response under nutritional stresses increases expression of virulence-associated genes encoded on Francisella Pathogenicity Island. This study investigated how stringent response governs the oxidative stress response of *F. tularensis*. We demonstrate that RelA/SpoT-mediated ppGpp production alters global gene transcriptional profile of *F. tularensis* in the presence of oxidative stress. The lack of stringent response in *relA/spoT* gene deletion mutants of *F. tularensis* makes bacteria more susceptible to oxidants, attenuates survival in macrophages, and virulence in mice. Mechanistically, we provide evidence that the stringent response in *Francisella* contributes to oxidative stress resistance by enhancing the production of antioxidant enzymes.

**Importance:** The unique intracellular life cycle of *Francisella* in addition to nutritional stress also exposes the bacteria to oxidative stress conditions upon its brief residence in the phagosomes, and escape into the cytosol where replication takes place. However, the contribution of the stringent response in gene regulation and management of the oxidative stress response when *Francisella* is experiencing oxidative stress conditions is not known. Our results provide a link between the stringent and oxidative stress responses. This study further improves our understanding of the intracellular survival mechanisms of *F. tularensis*.

## Introduction

*Francisella tularensis* is a Gram-negative bacterium responsible for causing tularemia in the northern hemisphere. *F. tularensis* has long been developed as a biological weapon due to its ability to cause severe illness upon inhalation of as few as ten organisms and based on its potential to be used as a bioterror agent is now classified as a Tier 1 Category A select agent by the CDC (1–3). The virulent strains are classified under *F. tularensis* subsp. *tularensis,* (type A) and *F. tularensis* subsp. *holarctica* (type B), whereas avirulent strains belong to *F*. *novicida* or *F. philomiragia* (4). The virulent SchuS4 strain belongs to *F. tularensis* subsp. *tularensis* while the live vaccine strain (LVS) is derived from *F. tularensis* subsp. *holarctica*. *Francisella* is a facultative intracellular pathogen and can replicate in a variety of cell types; however, macrophages are the primary sites of replication (5, 6). The clinical presentation of tularemia depends on the route, dose, and infecting strain of *F. tularensis*. The ulceroglandular, oculoglandular, or the typhoidal forms of tularemia are not fatal. However, pneumonic tularemia is a highly acute and fatal form of the disease.

The unique intracellular life cycle of *Francisella* exposes the bacteria to oxidative stress conditions upon its entry, brief residence in the phagosomes, and escape from phagosomes into the cytosol where replication takes place (5). *F. tularensis* genome encodes conventional antioxidant enzymes such as Fe- and CuZn-containing superoxide dismutases (SodB and SodC, respectively), catalase (KatG), and alkyl hydroperoxide reductase (AhpC) homologs to counter the oxidative stress generated at these distinct intracellular locations (7–10). Unlike other bacterial pathogens, SoxR, an oxidative stress response regulator which regulates the expression of SodB, SodC and SodA (manganese containing Sod) is absent in *F. tularensis*. Instead, another oxidative stress regulator OxyR is present, which regulates the expression of KatG and AhpC in *F. tularensis* (11). In *F. tularensis* SodB, KatG, and AhpC are induced in response to oxidative stress and are secreted in abundance in the extracellular milieu and into the cytosol of *F. tularensis* infected macrophages (12). Both the SodB and KatG are secreted by the Type I Secretion System of *F. tularensis* (13). Expression of these primary antioxidant genes starts immediately upon phagocytosis of *F. tularensis* SchuS4 and remains significantly upregulated during phagosomal and cytosolic phases suggesting that *F. tularensis* experiences oxidative stress at both of these intracellular locations (14).

The stringent response facilitates bacterial survival under nutritionally challenging starvation conditions. The hallmark of stringent response is the accumulation of the effector molecules ppGpp and (p)ppGpp (guanosine-5’-diphosphate-3’-diphosphate and guanosine-5’-triphosphate-3’-diphosphate) known as stress alarmones (15). The *relA* and *spoT* gene products generate alarmones in several Gram-negative bacterial pathogens. RelA is a ribosome-associated ppGpp synthetase that gets activated under amino acid starvation conditions whereas, SpoT is a bifunctional enzyme with both ppGpp synthetase and ppGpp hydrolase activities. During amino acid starvation, the presence of uncharged tRNA in the acceptor site of the ribosomes sends a signal to the RelA protein associated with ribosomes to catalyze the phosphorylation of GTP in conjunction with ATP as a donor to generate (p)ppGpp. The cytosolic SpoT is required for the basal synthesis of ppGpp during bacterial growth, (p)ppGpp degradation and elevated synthesis of ppGpp under several stress conditions, including fatty acid and carbon starvation (16). The ppGpp cause global reprogramming of cellular and metabolic function by binding to the β’-subunit of the RNA polymerase to activate or repress several genes or by interacting directly with proteins to promote adaptation, survival, and transmission in adverse growth conditions.

*Francisella* encodes a monofunctional *relA* and a bifunctional *spoT* gene. A single *relA* gene deletion mutant of *F. novicida* grows better than the wild type bacteria, but exhibits attenuated virulence in a mouse model of tularemia (17). *F. tularensis* utilizes *relA/spoT* mediated ppGpp production to promote stable physical interactions between the components of the transcription machinery to activate expression of virulence-associated genes encoded on Francisella Pathogenicity Island (FPI). It has also been shown that SpoT rather than RelA-dependent production of ppGpp is essential for the expression of virulence genes in *F. tularensis* (18). In addition to *relA/spoT*; *migR, trmE,* and *chpA* genes of *F. tularensis* through unidentified mechanisms are also involved in ppGpp production, and regulation of FPI encoded virulence-associated genes (19). A global gene expression profile of *F. tularensis* SchuS4 under nutrient limitation conditions induced by serine hydroxamate showed ppGpp-dependent upregulation of genes involved in virulence, metabolism and stress responses associated with downregulation of genes required for transport and cell division (20). These studies demonstrate that stringent response under environmental and nutritional stresses increase FPI gene expression. However, the contribution of the stringent response in gene regulation and management of the oxidative stress response when *Francisella* is experiencing oxidative stress conditions is not known.

In this study, we investigated how stringent response governs the oxidative stress response of *F. tularensis*. Our results provide a link between the stringent and oxidative stress responses. We demonstrate that *relA/spoT*-mediated ppGpp production alters global gene transcriptional profile of *F. tularensis* in the presence of oxidative stress. The lack of stringent response in *relA/spoT* gene deletion mutants of *F. tularensis* makes bacteria more susceptible to oxidants, attenuates survival in macrophages, and virulence in mice. Mechanistically, we provide evidence that the stringent response in *Francisella* contributes to oxidative stress resistance by enhancing the production of several antioxidant enzymes, including KatG and SodB. This study further enhances our understanding of the intracellular survival mechanisms of *F. tularensis*.

## Results

The *relA* gene (*FTL_0285*) encoding GTP Pyrophosphokinase in *Francisella* is 647 base pairs in length and is transcribed as a single transcriptional unit; while the *spoT* gene (*FTL_1413*) encoding a guanosine-3’ 5’-bis (diphosphate) 3’-phosphohydrolase/(p)ppGpp synthase is 704 bp in length is transcribed as an operon along with *capA, capB* and *capC* genes (Fig. 1A). To investigate the role of stringent response in oxidative stress resistance of *F. tularensis* LVS, we constructed unmarked, in-frame *relA* single gene deletion (Δ*relA*) and *relA spoT* (Δ*relA*Δ*spoT*) double gene deletion mutants. The gene deletions in single and double mutants were confirmed by PCR (Fig. 1B) followed by DNA sequencing. The transcomplemented strains were constructed by providing full copies of the *relA* gene in the Δ*relA* mutant, and *spoT* gene in the Δ*relA*Δ*spoT* mutant. To determine the ability of the wild type *F. tularensis* LVS, Δ*relA* and the Δ*relA*Δ*spoT* mutants to produce ppGpp, the bacterial strains were grown to late exponential phase, and ppGpp production was determined by HPLC. The production of ppGpp was dropped to 45 and 28% in the Δ*relA* and the Δ*relA*Δ*spoT* mutants respectively, as compared to that observed for the wild type *F. tularensis* LVS, indicating that Δ*relA* and more so the Δ*relA*Δ*spoT* mutant is deficient in ppGpp production (Fig. 1C and D). The growth characteristics of the wild type *F. tularensis* LVS, the Δ*relA,* the Δ*relA*Δ*spoT* double mutant and the transcomplemented strains were determined by growing in MH-broth in the absence or presence of 500µM of serine hydroxamate, a serine homolog responsible for inducing amino acid starvation-like conditions. The Δ*relA* mutant did not exhibit any growth defect and grew similarly to the wild type *F. tularensis* LVS when grown in the MH-broth. The Δ*relA*Δ*spoT* mutant showed retarded growth and entered the stationary phase after 16 hours of growth. Transcomplementation with *spoT* gene restored the growth of the Δ*relA*Δ*spoT* mutant as well as prevented its early entry into the stationary phase (Fig 1E). Growth of all *Francisella* strains was slightly reduced in the presence of serine hydroxamate; however, the Δ*relA*Δ*spoT* mutant failed to grow in its presence. Transcomplementation of the Δ*relA*Δ*spoT* with the *spoT* gene restored its growth in the presence of serine hydroxamate (Fig. 1F). Sensitivities of the wild type *F. tularensis* LVS, the Δ*relA*, the Δ*relA*Δ*spoT* mutants and the transcomplemented strains were also tested against streptomycin, nitrofurantoin, and tetracycline by disc diffusion assays. It was observed that the Δ*relA*Δ*spoT* mutant exhibited enhanced sensitivities towards all the three antibiotics tested as compared to the wild type *F. tularensis* LVS or the Δ*relA* mutant. Transcomplementation of the Δ*relA*Δ*spoT* mutant restored the sensitivities similar to those observed for the wild type or the Δ*relA* mutant (Fig. 1G). Collectively, these results demonstrate that induction of RelA/SpoT-mediated stringent response associated with the production of ppGpp is required for growth of *F. tularensis* under normal growth conditions, amino acid starvation, as well as resistance towards antibiotics.

**Figure 1.**
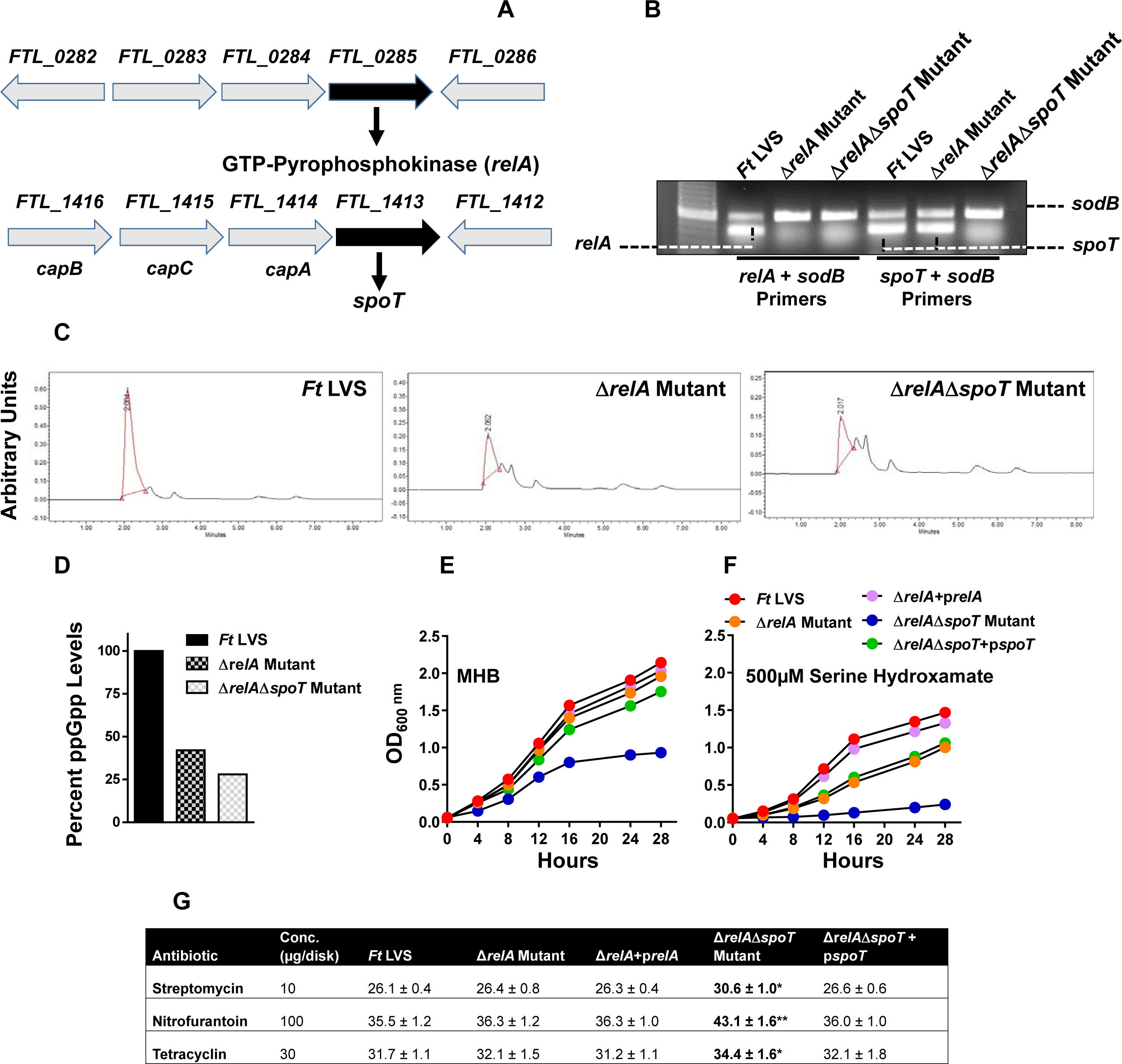
Genomic organization, generation, and characterization of the relA and relA/spoT gene deletion mutants of the F. tularensis LVS. (A) The genomic organization of the *relA* and the *spoT* genes of the *F. tularensis* LVS. (B) Multiplex colony PCR using *relA* and *spoT* gene-specific primers and *sodB* gene primers as internal controls. Amplification of the *sodB* gene confirmed the presence of the DNA template in the reaction, whereas the absence of the *relA* gene product in the Δ*relA* and Δ*relA*Δ*spoT* mutants confirmed the deletion of the *relA* gene and that of *spoT* confirmed *spoT* gene deletion in the Δ*relA*Δ*spoT* mutant. (C and D) Determination of ppGpp production and its quantitation in the *F. tularensis* (*Ft*) LVS, the Δ*relA* and the Δ*relA*Δ*spoT* mutants by HPLC. (E and F) Growth curves of the *Ft* LVS, the Δ*relA*, the Δ*relA*Δ*spoT* mutants, and the transcomplemented strains in the presence of the indicated concentrations of serine hydroxamate. (G) The susceptibility of the *Ft* LVS, the Δ*relA* and the Δ*relA*Δ*spoT* mutants and the transcomplemented strains to indicated antibiotics was tested by disk diffusion assay. The plates were incubated for 48-72 hours, and the zones of inhibition around the discs were measured. The results shown in C, D, E, F, and G are representative of three independent experiments with identical results. The data in G are represented as Mean±S.D. (n=3 biological replicates) and were analyzed using one-way ANOVA (\**P*<0.05; \*\**P*<0.01).

### Transcriptional profile of the Δ*relA*Δ*spoT* mutant

RNA sequencing was used to obtain the transcriptional profiles of the wild type *F. tularensis* LVS and the Δ*relA*Δ*spoT* mutant with or without the treatment with H_2_O_2_. Since the Δ*relA*Δ*spoT* mutant grow slower than the wild type *F. tularensis* LVS in the presence of H_2_O_2,_ 3-hour treatment with H_2_O_2_ was selected. At this time point post-treatment, the viability of both the wild type and the mutant strain is identical. A *P* value of *<0.05* was used as a cut-off to verify differentially expressed genes in the Δ*relA*Δ*spoT* mutant. A total of 318 genes were differentially expressed in untreated Δ*relA*Δ*spoT* mutant as compared to the wild type *F. tularensis* LVS. Of these 163 genes (51%) were downregulated while 155 genes (49%) were upregulated in the Δ*relA*Δ*spoT* mutant. Majority of the downregulated genes were involved in metabolism (n=63), hypothetical proteins (n=30), Francisella Pathogenicity Island (FPI) genes (n=30) as well as transport, replication, transcription, translation and stress response genes. The upregulated genes mostly belonged to the hypothetical proteins (n=65) metabolism (n=47) and translation (n=12) categories (Fig. 2). Treatment of the Δ*relA*Δ*spoT* mutant with H_2_O_2_ resulted in differential expression of a total of 855 genes in the Δ*relA*Δ*spoT* mutant as compared to the wild type *F. tularensis* LVS. Of the 450 (53%) downregulated genes, majority of the genes belonged to metabolism (n=155), hypothetical proteins (n=95), others (n=49) and FPI (n=32) categories. The genes that were upregulated following exposure to H_2_O_2_ also belonged to the similar categories except for the FPI genes (Fig. 2). Collectively these results demonstrate that RelA and SpoT not only control the expression of several genes of *F. tularensis* under normal growth conditions but also regulate genes both positively and negatively when the bacteria are exposed to the oxidative stress.

**Figure 2.**
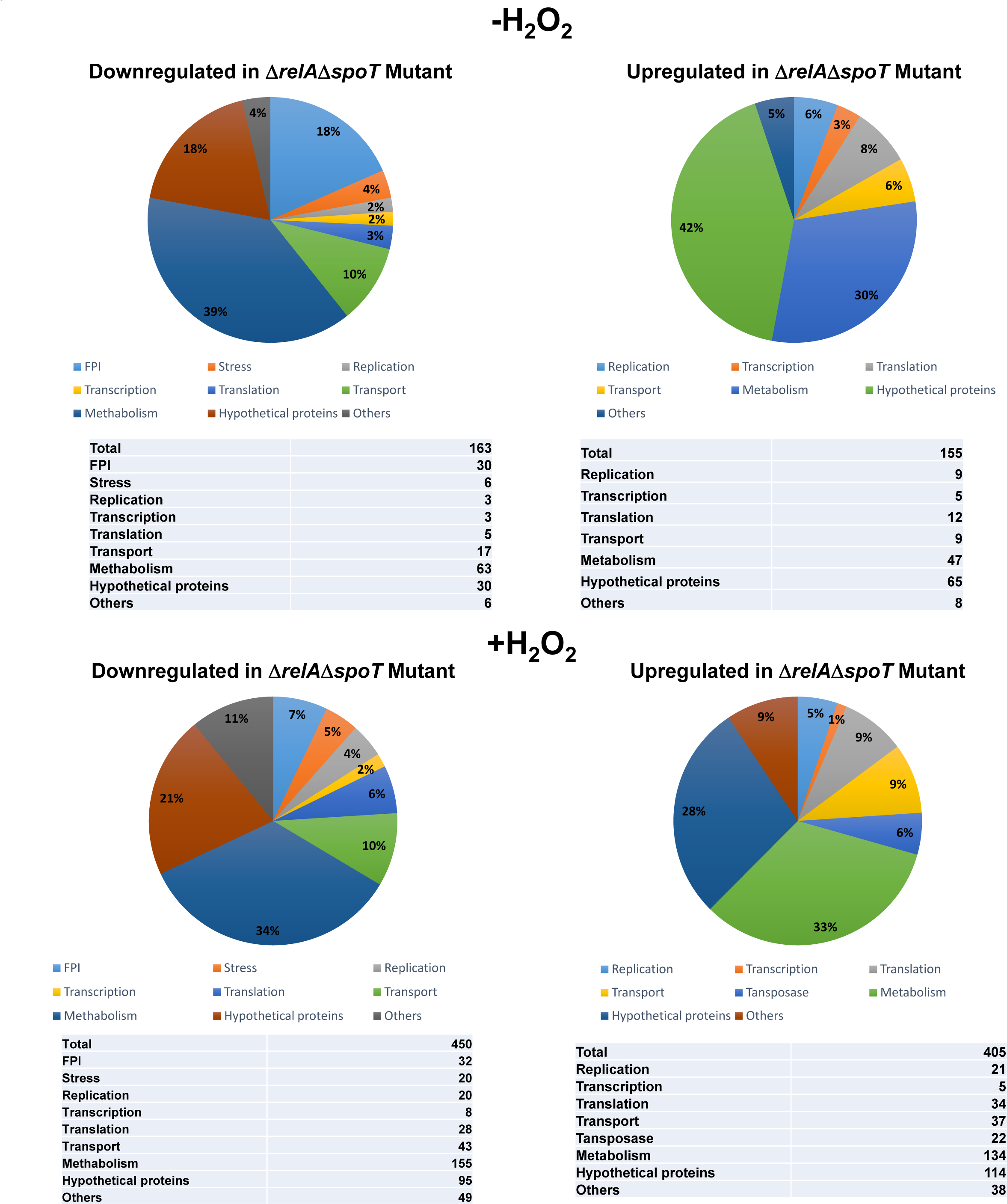
Differential expression of genes in the ΔrelAΔspoT mutant of F. tularensis with or without exposure to H_2_O_2_. (A) A total number of differentially expressed genes in the Δ*relA*Δ*spoT* mutant compared to the wild-type *F. tularensis* LVS.

### Expression of the FPI genes

We further analyzed the differential expression of the FPI genes in the Δ*relA*Δ*spoT* mutant under normal as well as oxidative stress conditions created by exposing the bacteria to H_2_O_2._ The genes encoded on FPI were significantly downregulated in untreated Δ*relA*Δ*spoT* mutant as compared to the wild type *F. tularensis* LVS. The expression of *iglA, B, C,* and *D* genes remained similarly downregulated when the Δ*relA*Δ*spoT* mutant was exposed to H_2_O_2._ Contrary to the *igl* locus, the expression of *pdpC, iglJ, IglI,* and *dotU* genes which are encoded on another FPI operon was significantly downregulated (*P adjusted value <0.05*) when the Δ*relA*Δ*spoT* mutant was exposed to H_2_O_2_ than the untreated Δ*relA*Δ*spoT* mutant bacteria as compared to treated or untreated wild type *F. tularensis* LVS. The expression of *iglF, vgrG, iglE, pdpB,* and *pdpA* which are encoded on a separate FPI operon was slightly repressed with maximum downregulation observed for the *vgrG* gene (*P adjusted value <0.001*) when the Δ*relA*Δ*spoT* mutant bacteria were exposed to H_2_O_2._ These results indicate that stringent response induced by RelA/SpoT positively regulates the expression of virulence-associated genes encoded on the FPI under normal as well as oxidative stress conditions (Fig. 3A).

**Figure 3.**
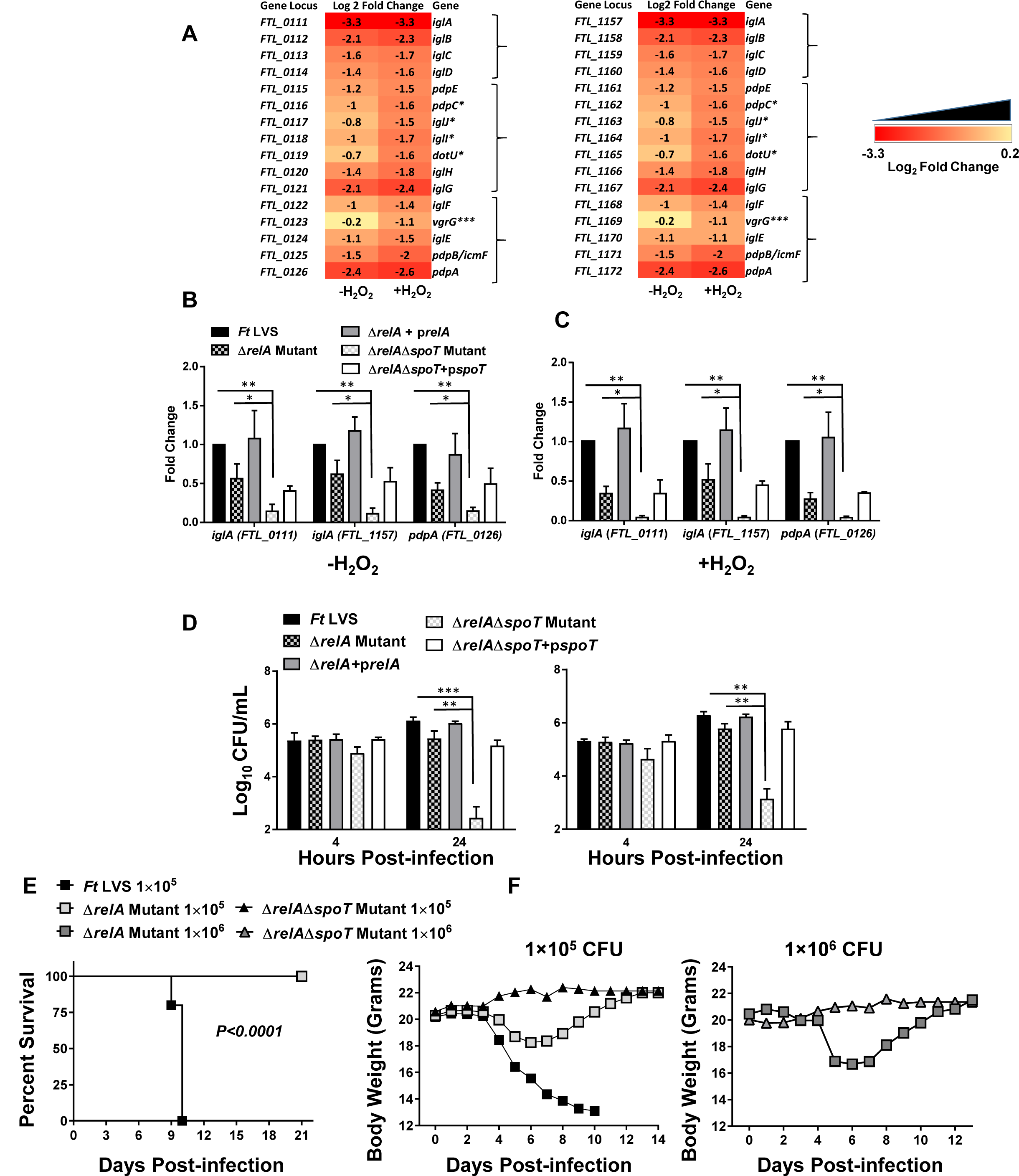
Differential expression of genes encoded on FPI in the absence or the presence of oxidative stress, intramacrophage survival and virulence of ΔrelA and ΔrelAΔspoT mutants of F. tularensis. (A) RNAseq analysis comparing the downregulation of important virulence genes in the Δ*relA*Δ*spoT* mutant compared to that of wild type *Ft* LVS. The data are represented as Log_2_ fold change and are cumulative of three independent experiments. The brackets on the right represent operon. (B and C) Quantitative reverse transcriptase PCR (qRT-PCR) was performed to evaluate the transcription of select genes. The amount of target gene amplification was normalized to a *tul4* internal control. The relative mRNA levels are presented as Mean±SD (n=3 biological replicates). (D) RAW macrophage cell line (left panel) and Bone Marrow-Derived Macrophages (BMDMs) (right panel) were infected with wild type *F. tularensis* (*Ft*) LVS, the indicated mutants and the transcomplemented strains at a multiplicity of infection (MOI) of 100. The cells were lysed 4 and 24 hours post-infection, diluted 10-fold, and plated for enumeration of bacterial colonies. The data are represented as Mean±SD (n=3 biological replicates) and are a representative of 4 independent experiments. (E and F) C57BL/6 mice (n=5/group) were infected intranasally with 1×10^5^ CFUs of *F. tularensis* LVS and 1×10^5^ or 1×10^6^ CFUs of the Δ*relA* and the Δ*relA*Δ*spoT* mutants. The mice were observed for mortality and morbidity for 21 days. The body weights of the infected mice were recorded daily to monitor the progress of infection. The data in B, C, and D were analyzed using ANOVA and E using the Log-rank test. (\**P*<0.05, \*\**P*<0.01, \*\*\**P*<0.001).

We also confirmed the expression profiles of select FPI genes in the wild type *F. tularensis* LVS, the Δ*relA* and the Δ*relA*Δ*spoT* mutants by quantitative real-time PCR (qRT-PCR). It was observed that deletion of the *relA* gene affected the expression of *iglA* genes encoded on both copies of the FPI as well as the *pdpA* gene. However, these genes are significantly downregulated in the Δ*relA*Δ*spoT* mutant as compared to the wild type *F. tularensis* LVS and the Δ*relA* mutant both in untreated, as well as H_2_O_2_ treated bacteria. Transcomplementation of the Δ*relA* mutant completely restored the expression levels of the FPI genes, while the expression of these genes was only partially restored in the Δ*relA*Δ*spoT* mutant transcomplemented with the *spoT* gene (Fig. 3B and C).

We next performed cell culture assays using the Raw 264.7 cell line and BMDMs to investigate if the downregulated expression of the FPI genes is associated with attenuated intramacrophage survival. Except the Δ*relA*Δ*spoT* mutant, equal numbers of bacteria were taken up by the infected Raw cells or BMDMs after 4 hours post-infection. Nearly 5-fold fewer bacteria were taken up by the macrophages infected with the Δ*relA*Δ*spoT* mutant at 4-hour time-point. At 24 hours post-infection, the Δ*relA* mutant was found to be only partially attenuated for intramacrophage growth. However, nearly 20-fold less Δ*relA*Δ*spoT* mutant bacteria were recovered at 24 hours post-infection as compared to the wild type *F. tularensis* indicating an attenuation of the intramacrophage growth. Transcomplementation of both the Δ*relA* and Δ*relA*Δ*spoT* mutants restored the intramacrophage replication (Fig. 3D).

We also investigated if the loss of *relA* and *relA spoT* is associated with the attenuation of virulence in mice. It was observed that 100% of C57BL/6 mice infected intranasally either with 1×10^5^ or 1×10^6^ CFUs of the Δ*relA* or the Δ*relA*Δ*spoT* mutant survived the infection. Mice infected with the Δ*relA* mutant exhibited initial loss of body weight from days 4-8 post-infection and then regained their initial body weight. On the other hand, mice infected with the Δ*relA*Δ*spoT* mutant did not show any body weight loss. All the control mice infected with 1×10^5^ CFU of *F. tularensis* LVS succumbed to infection by day ten post-infection and experienced severe body weight loss (Fig. 3E and F). Collectively, these results demonstrate that RelA and SpoT are required for expression of virulence-associated genes encoded on FPI, intramacrophage survival, and virulence in mice.

### Expression of stress and heat shock proteins

Induction of stress-related proteins is a hallmark of the RelA-SpoT-dependent stringent response. We investigated the expression profiles of genes encoding key stress-related proteins in the Δ*relA*Δ*spoT* mutant in the absence or the presence of oxidative stress and compared those with the wild type *F. tularensis* LVS. Three distinct gene expression profiles were observed. The expression of *clpB, hsp90,* and *hsp40* genes remained unaltered in the Δ*relA*Δ*spoT* mutant when compared with the wild type *F. tularensis* LVS in the absence of oxidative stress. However, these genes were found to be downregulated in the Δ*relA*Δ*spoT* mutant as compared to the wild type *F. tularensis* LVS when exposed to the oxidative stress. All these three genes are transcribed as single transcription units. Genes encoding Clp ATPases *clpP, clpX,* and *lon*; *hslU, hslV; grpE, dnaK,* and *dnaJ* were downregulated in untreated Δ*relA*Δ*spoT* mutant and were further downregulated when exposed to the oxidative stress induced by H_2_O_2._ A group of genes encoding the universal stress protein, protease *sohB*, and starvation protein A were down-regulated in untreated Δ*relA*Δ*spoT* mutant and remained downregulated upon exposure to the oxidative stress (Fig. 4A). The differential expression of these genes was also confirmed by qRT-PCR. The expression profile of the representative stress response genes was similar to that observed by RNAseq. However, all these genes were significantly downregulated in untreated as well as H_2_O_2_ treated Δ*relA*Δ*spoT* mutant as compared to the *F. tularensis* LVS. Transcomplementation of the Δ*relA* mutant restored the wild type phenotype, while only a partial restoration of the gene expression was observed in the Δ*relA*Δ*spoT* mutant (Fig. 4B).

**Figure 4.**
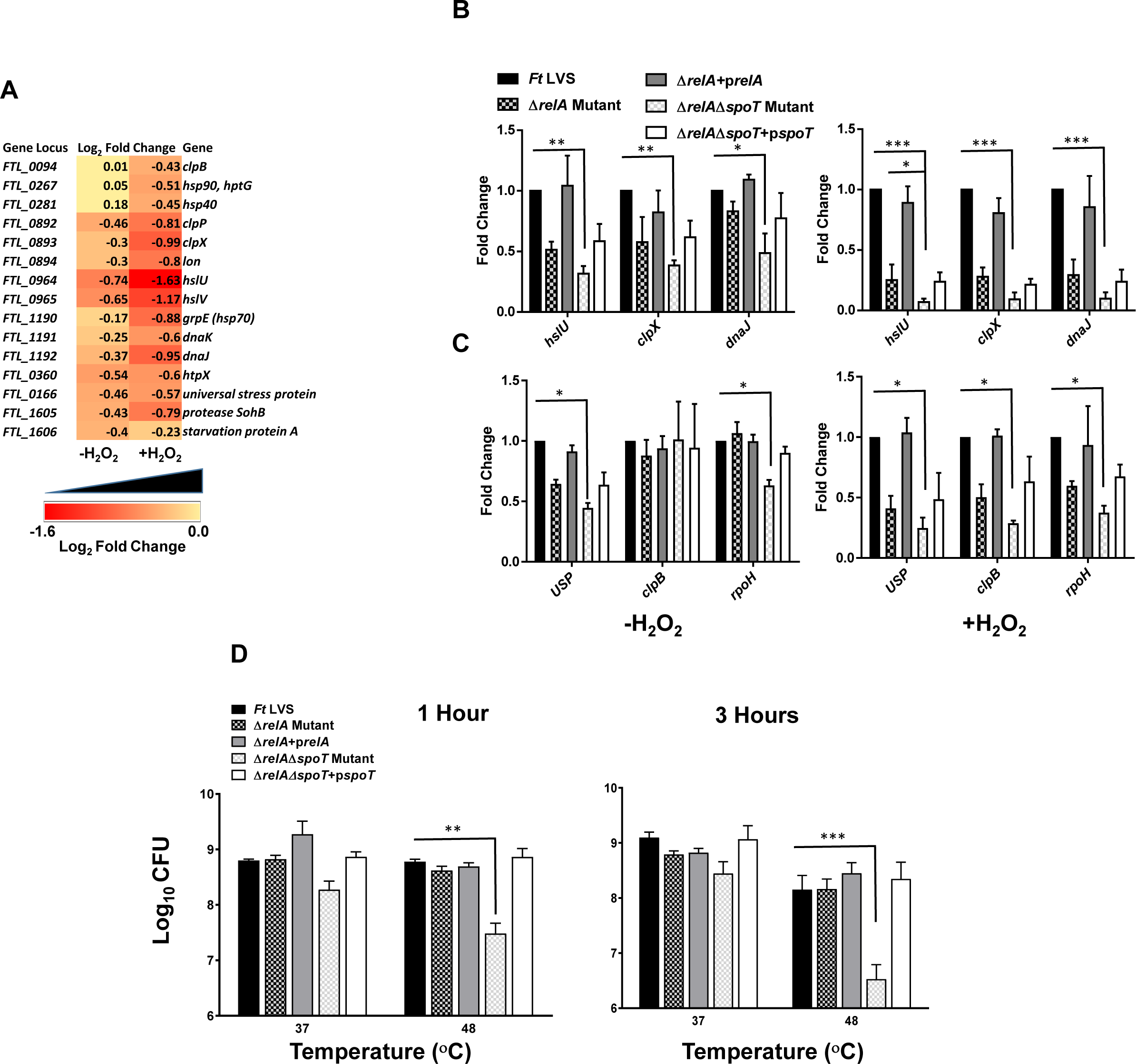
Differential expression of stress response genes in the absence or the presence of oxidative stress and sensitivity of the ΔrelA and ΔrelAΔspoT mutants of F. tularensis to a higher temperature. (A) RNAseq analysis comparing the downregulation of important stress response genes in the Δ*relA*Δ*spoT* mutant compared to that of the wild type *Ft* LVS. The data are represented as Log_2_ fold change and are cumulative of three independent experiments. (B and C) Quantitative reverse transcriptase PCR (qRT-PCR) was performed to evaluate the transcription of select genes. The amount of target gene amplification was normalized to a Tul4 internal control. The relative mRNA levels are presented as Mean±SD (n=3 biological replicates). (D) A cell viability assay was performed by growing the indicated bacterial strains at 37 and 48^ο^C for 1 and 3 hours. The cultures were diluted 10-fold and spotted on MH-chocolate agar plates. The data in C and D were analyzed by ANOVA, and the *P* values were recorded. \**P*<0.05; \*\**P*<0.01; \*\*\**P*<0.001.

Since RelA-SpoT-dependent induction of heat shock proteins is associated with survival at higher temperatures, we also examined the effect of the loss of *relA* and *relA/spoT* on bacterial viability when exposed to a higher temperature of 48^ο^C. It was observed that the viability of the Δ*relA* mutant was unaffected and remained similar to the wild type *F. tularensis* when exposed to a temperature of 48^ο^C for 1 hour and 5-7 fold fewer bacteria were recovered after 3 hours of exposure. On the other hand, nearly 10-20-fold less viable bacteria were recovered when the Δ*relA*Δ*spoT* mutant was exposed to a temperature of 48^ο^C for 1 and 3 hours, respectively. Transcomplementation of the Δ*relA*Δ*spoT* mutant restored the wild type phenotype (Fig. 4D). Collectively, these results indicate that RelA-SpoT-mediated stringent response induces protective mechanisms by altering the expression of genes involved in degradation and disaggregation accumulated or misfolded proteins, and DNA damage repair mechanisms under the conditions of oxidative stress. Also, RelA-SpoT-mediated stringent response facilitates bacterial survival at higher temperatures.

### Expression of MglA-dependent genes and transcriptional regulators

It has been reported the ppGpp produced by RelA and SpoT plays an important role in MglA-dependent gene regulation of *F. tularensis* (18). We next investigated if the transcription of the MglA-dependent genes is altered in untreated or H_2_O_2_ treated Δ*relA*Δ*spoT* mutant. It was observed that all the major MglA-regulated genes remained similarly downregulated in untreated as well as H_2_O_2_ treated Δ*relA*Δ*spoT* mutant. These results were also confirmed by qRT-PCR of the highly downregulated MglA-dependent gene *FTL_1219* (Fig. 5A and 5B).

**Figure 5.**
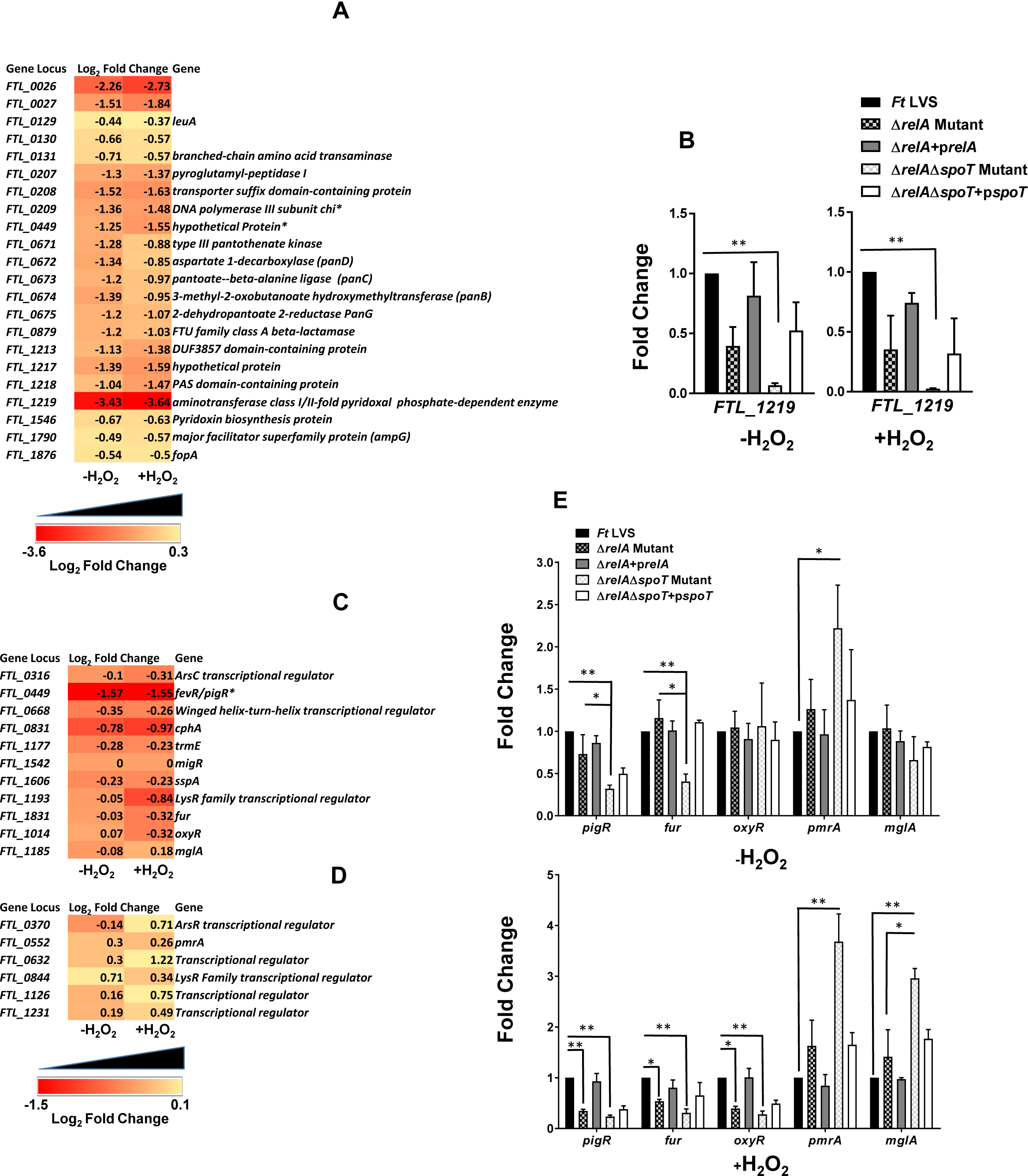
Differential expression of MglA regulated and transcriptional regulator genes in the absence or the presence of oxidative stress. (A) RNAseq analysis comparing the downregulation of important MglA regulated genes in the Δ*relA*Δ*spoT* mutant compared to that of the wild type *Ft* LVS. (B) qRT-PCR was performed to evaluate the transcription of a select gene. (C and D) RNAseq analysis comparing the downregulation of important transcriptional regulator genes in the Δ*relA*Δ*spoT* mutant compared to that of the wild type *Ft* LVS. The data are represented as Log_2_ fold change and are cumulative of three independent experiments. (E) qRT-PCR analysis to evaluate the transcription of select genes. The data shown in A, C, and D are represented as Log_2_ fold change and are cumulative of three independent experiments. In B and E, the amount of target gene amplification was normalized to a *tul4* internal control. The relative mRNA levels are presented as Mean±SD (n=3 biological replicates). The data in B and E were analyzed by ANOVA, and the *P* values were recorded. \**P*<0.05; \*\**P*<0.01.

We also investigated if the loss of stringent response mediated by RelA/SpoT affects the expression of other transcription regulators in the absence or the presence of oxidative stress. It was observed that the expression of the majority of the transcriptional regulators was downregulated in untreated as well as H_2_O_2_ treated Δ*relA*Δ*spoT* mutant. Transcription of *fevR/pigR* and *cphA* was maximally downregulated in the Δ*relA*Δ*spoT* mutant under both the conditions tested. The expression of *migR* remained unaltered and was found to be similar to that observed for the wild type *F. tularensis,* while the expressions of LysR family transcriptional regulator (*FTL_1831*), *fur* and *oxyR* were downregulated upon exposure to H_2_O_2_ (Fig. 5C). On the other hand, transcript levels of several regulators including *pmrA*, *FTL_0632, FTL_ 1126* and *mglA* were upregulated in the H_2_O_2_ treated Δ*relA*Δ*spoT* mutant (Fig. 5D). Comparison of the differential expression of *pigR, fur, pmrA* and the *mglA* levels between the untreated and H_2_O_2_ treated Δ*relA* and the Δ*relA*Δ*spoT* mutants by qRT-PCR revealed that the expression of these genes was not affected in untreated Δ*relA* mutant and remained similar to the wild type *F. tularensis* LVS. On the other hand, the expression of *pigR* and *fur* was significantly downregulated as compared to *F. tularensis* LVS and the Δ*relA* mutant, and that of *pmrA* was upregulated in the untreated Δ*relA*Δ*spoT* mutant as compared to *F. tularensis* LVS. Transcript levels of *oxyR* for all three untreated bacterial strains tested remained unaltered. However, exposure to H_2_O_2_ downregulated the expression of the *pigR, Fur and oxyR* in the Δ*relA* and Δ*relA*Δ*spoT* mutants from the levels observed in their untreated counterparts. The transcript levels of both the *pmrA* and *mglA* were upregulated in H_2_O_2_ treated Δ*relA*Δ*spoT* mutant. Transcomplementation of both the mutants restored the wild type phenotype (Fig. 5E). Taken together, these results demonstrate that RelA-SpoT mediates the differential expression of several transcriptional regulators under the normal homeostatic as well as the conditions of oxidative stress.

### Expression of genes involved in antioxidant defense mechanisms

We next investigated the effect of loss of stringent response in the expression of genes encoding antioxidant enzymes of *F. tularensis*. Expression of *methionine sulfoxide reductase B* (*msrB*)*, thioredoxin* and *thioredoxin 1* (*FTL_0611* and *FTL_1224*) *glutaredoxin 3 and 1*, *glutathione peroxidase,* and *hypothetical protein* genes co-transcribed with *msrB* and *thioredoxin 1* (*trx1*) were downregulated in the Δ*relA*Δ*spoT* mutant and remained similarly downregulated following its exposure to H_2_O_2_. The transcript levels of *short chain dehydrogenase*, *katG,* and *sodB* genes were marginally downregulated in untreated Δ*relA*Δ*spoT* mutant. The expression of the *short-chain dehydrogenase* gene was upregulated, while those of *katG* and *sodB* remained unaltered following treatment of the Δ*relA*Δ*spoT* mutant with H_2_O_2_. The expression of the other two primary antioxidant enzymes *sodC* and *ahpC* remained unaltered in the untreated Δ*relA*Δ*spoT* mutant while their transcript levels were upregulated in the H_2_O_2_ treated Δ*relA*Δ*spoT* mutant (Fig. 6A). The expression levels of *sodB, katG, thioredoxin 1,* and *glutaredoxin 2* were also confirmed by qRT-PCR, and the results similar to those obtained from the RNAseq were observed (Fig. 6B). We also tested the protein levels of two primary antioxidant enzymes KatG and SodB in bacterial lysates from the wild type *F. tularensis* LVS, the Δ*relA*, Δ*relA*Δ*spoT* mutant and the transcomplemented strains by western blot analysis using anti-KatG and anti-SodB antibodies. Diminished protein levels of both KatG and SodB were observed in the Δ*relA* mutant and more so in the Δ*relA*Δ*spoT* mutant and these levels were marginally improved in the transcomplemented strains (Fig. 6C). Collectively, these results indicate that the loss of stringent response is associated with differential expression of genes involved in antioxidant defense mechanisms of *F. tularensis*.

**Figure 6.**
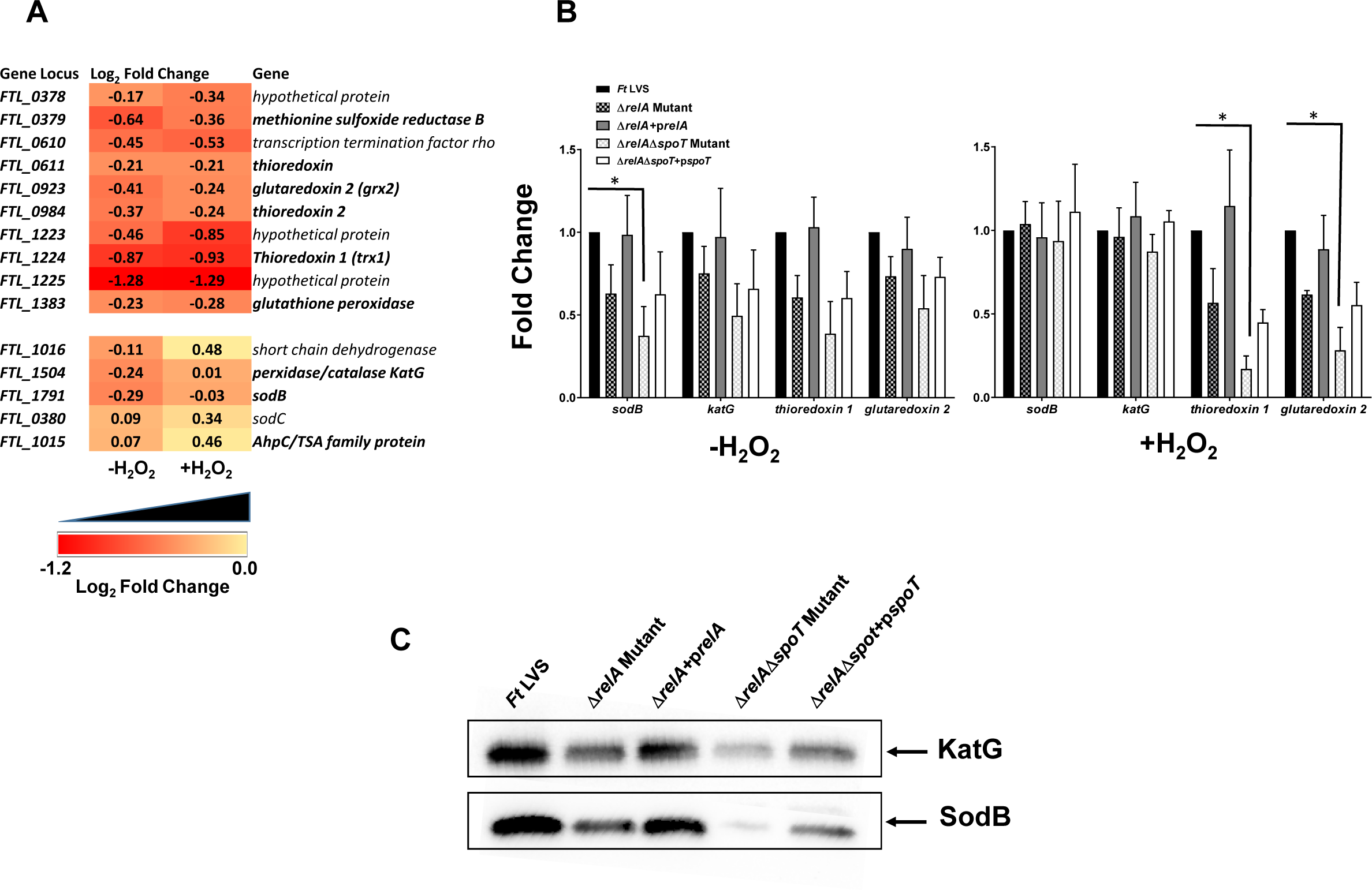
Differential expression of antioxidant enzyme genes in the absence or the presence of oxidative stress. (A) RNAseq analysis comparing the downregulation of important antioxidant enzyme genes in the Δ*relA*Δ*spoT* mutant compared to that of the wild type *Ft* LVS. (B) qRT-PCR was performed to evaluate the transcription of a select gene. The data shown in A are represented as Log_2_ fold change and are cumulative of three independent experiments. In B, the amount of target gene amplification was normalized to a *tul4* internal control. The relative mRNA levels are presented as Mean±SD (n=3 biological replicates). The data in B and E were analyzed by ANOVA, and the *P* values were recorded. \**P*<0.05. (C) The western blots of the lysates of the indicated *Francisella* strains probed with anti-KatG antibodies, that were stripped and re-probed with antibodies against SodB.

### Δ*relA*Δ*spoT* mutant of *F. tularensis* LVS is hypersensitive to oxidative stress

We next investigated the sensitivity of the Δ*relA*Δ*spoT* mutant by performing growth curve analysis, bacterial killing, and disc diffusion assays in the presence of oxidants. As observed earlier, the Δ*relA*Δ*spoT* mutant grew slowly and entered the stationary phase earlier than the wild type *F. tularensis* LVS or the Δ*relA* mutant. Transcomplementation of the Δ*relA*Δ*spoT* mutant restored its growth (Fig. 7A). However, the Δ*relA*Δ*spoT* mutant failed to grow when the growth curves were generated in the presence of 0.5 and 1.0 mM H_2_O_2_ and again, transcomplementation of the Δ*relA*Δ*spoT* mutant restored its growth similar to that observed for the Δ*relA* mutant (Fig. 7B). We confirmed these findings by performing bacterial killing assays by exposing wild type *F. tularensis,* the Δ*relA* and the Δ*relA*Δ*spoT* mutants and the transcomplemented strains to H_2_O_2_ for 1 and 3 hours and counting the colonies for bacterial viability. The viability of both the Δ*relA* and the Δ*relA*Δ*spoT* mutant was similarly affected after 1 hour of treatment. However, the viability of the Δ*relA*Δ*spoT* mutant was further reduced after 3 hours of exposure to H_2_O_2_ and nearly 30- and 20-fold less viable Δ*relA*Δ*spoT* mutant bacteria, respectively, were recovered as compared to those for the wild type *F. tularensis* LVS and the Δ*relA* mutant (Fig. 7C). Similar results were obtained when assays were conducted using the superoxide-generating compounds paraquat and pyrogallol (Fig. 7D and E). The Δ*relA*Δ*spoT* mutant also exhibited enhanced sensitivity as compared to the wild type *F. tularensis* LVS or the Δ*relA* mutant towards another superoxide-generating compound menadione and organic peroxide TBH when tested by disc diffusion assays (Fig. 7F and G). Collectively, these results demonstrate that the loss of *relA* and *spoT* is associated with enhanced susceptibility of the Δ*relA*Δ*spoT* mutant towards oxidative stress. These results also indicate that the stringent response induced by the RelA and SpoT is also linked with the oxidative stress resistance of *F. tularensis*.

**Figure 7.**
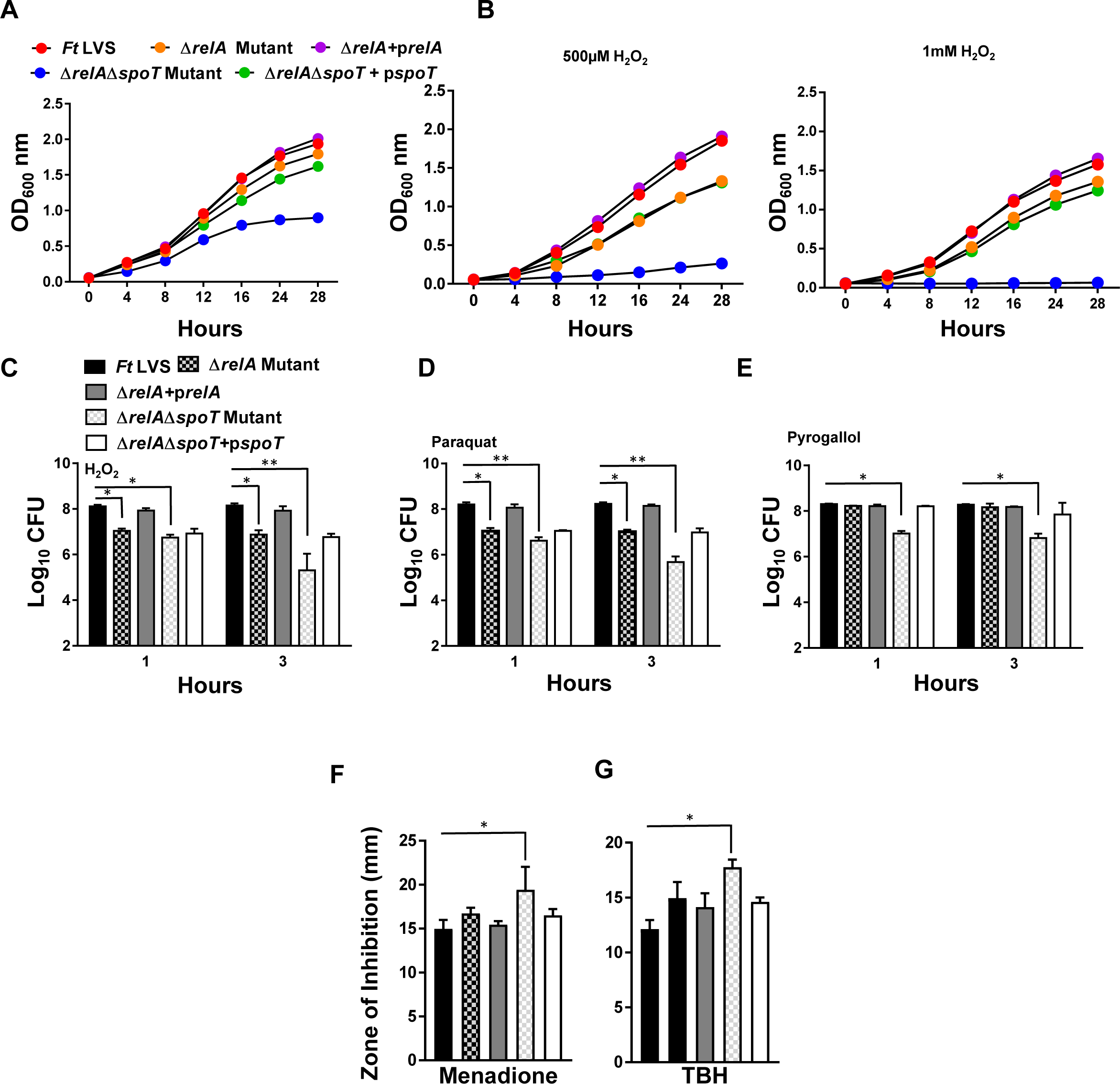
The sensitivity of the ΔrelA and ΔrelAΔspoT mutants of F. tularensis to oxidants. *F. tularensis* (*Ft*) LVS, *the* Δ*relA and* Δ*relA*Δ*spoT mutants* and the transcomplemented strains were grown in (A) MH-broth or (B) MH-broth containing 500 µM and 1mM of H_2_O_2_. The cultures were grown for 28 hours, and OD_600_ readings were recorded every 4 hours. The bacterial killing assay. The bacterial cultures were exposed to (C) 1000 µM of H_2_O_2,_ (D) 1mM Paraquat or (E)1mM Pyrogallol for 1 and 3 hours (n=3 biological replicates). The cultures were diluted 10- fold and plated on MH-chocolate agar plates for bacterial enumeration. The results are expressed as Log_10_ CFU/ml. The data are representative of three independent experiments conducted with identical results. Disc diffusion assay with superoxide-generating compounds, (F) menadione and (G) organic peroxide tert-butyl hydroperoxide (TBH). The data in C, D, E, F, and G were analyzed by ANOVA, and the *P* values were recorded. \**P*<0.05; \*\**P*<0.01.

### Overexpression of KatG and SodB restores the growth of Δ*relA*Δ*spoT* mutant of *F. tularensis* LVS

We observed that the Δ*relA*Δ*spoT* mutant revealed lower levels of primary antioxidant enzymes KatG and SodB and an enhanced sensitivity towards the oxidants. We investigated if overexpression of these two enzymes in the Δ*relA*Δ*spoT* mutant restores its resistance to oxidants. The Δ*relA*Δ*spoT* mutant was transcomplemented with *katG* (Δ*relA*Δ*spoT*+p*katG*) or the *sodB* gene (Δ*relA*Δ*spoT*+p*sodB*). Western blot analysis of the Δ*relA*Δ*spoT*+*pkatG* or Δ*relA*Δ*spoT*+p*sodB* revealed that protein levels of KatG and SodB were higher in Δ*relA*Δ*spoT*+p*katG* and Δ*relA*Δ*spoT*+p*sodB* strains, respectively, than those observed for the Δ*relA*Δ*spoT* mutant or the Δ*relA*Δ*spoT* mutant transcomplemented with the *spoT* gene (Fig. 8A). We next investigated the sensitivity of the Δ*relA*Δ*spoT*+p*katG* and Δ*relA*Δ*spoT*+p*sodB* strains towards oxidants. Expression of KatG or SodB in the Δ*relA*Δ*spoT* mutant did not alter the growth characteristics and the strains transcomplemented either with the *katG* or the *sodB* genes grew similar to the Δ*relA*Δ*spoT* mutant; while as observed earlier, transcomplementation with *spoT* partially restored the growth of the Δ*relA*Δ*spoT* mutant when grown in MH-broth (Fig. 8B and E). The growth of both the Δ*relA*Δ*spoT*+p*katG* and Δ*relA*Δ*spoT*+p*sodB* strains was restored partially to the Δ*relA*Δ*spoT*+p*spoT* levels when grown in the presence of H_2_O_2_ (Fig. 8C and F). As observed earlier, the Δ*relA*Δ*spoT* mutant failed to grow in the presence of H_2_O_2_. However, the growth of Δ*relA*Δ*spoT*+p*katG* or the Δ*relA*Δ*spoT*+p*sodB* strain was not restored when grown in the presence of serine hydroxamate and remained similar to that of the Δ*relA*Δ*spoT* mutant (Fig. 8D and G). We further investigated if the wild type phenotype is restored in the Δ*relA*Δ*spoT*+*pkatG* and the Δ*relA*Δ*spoT*+p*sodB* strains when exposed to superoxide-generating compounds and organic peroxides. It was observed that exposure of the Δ*relA*Δ*spoT*+*pkatG* strain to paraquat and TBH restored the wild type phenotype; while only a partial restoration was observed against menadione (Fig. 9A, B and C). On the other hand the wild type phenotype of the Δ*relA*Δ*spoT*+p*sodB* strain was restored only to menadione, but not when exposed either to paraquat or TBH (Fig. 9D, E and F). These results indicate that both KatG and SodB may be required for full restoration of the wild type phenotype in overcoming the oxidative stresses. It has previously been reported that stringent response mediates antibiotic resistance by modulating the pro- and antioxidant defenses. Our results showed that the Δ*relA*Δ*spoT* exhibit enhanced sensitivity towards antibiotics, we next investigated if this enhanced susceptibility is due to the oxidative stress and if overexpression of KatG or SodB reverts the antibiotic sensitivity of the Δ*relA*Δ*spoT* mutant. Our results show that antibiotic sensitivity of the Δ*relA*Δ*spoT*+*pkatG* and the Δ*relA*Δ*spoT*+p*sodB* strain remains similar to that observed for the Δ*relA*Δ*spoT* mutant (Fig. 9G) indicating that enhanced antibiotic sensitivity of the Δ*relA*Δ*spoT* mutant is independent of the oxidative stress mechanism. Collectively, these results demonstrate that loss of stringent response impairs the antioxidant defense mechanisms, thereby enhance the sensitivity of the Δ*relA*Δ*spoT* towards oxidants. However, overexpression of the primary antioxidant enzymes KatG and SodB restore the wild type phenotype associated with oxidant resistance.

**Figure 8.**
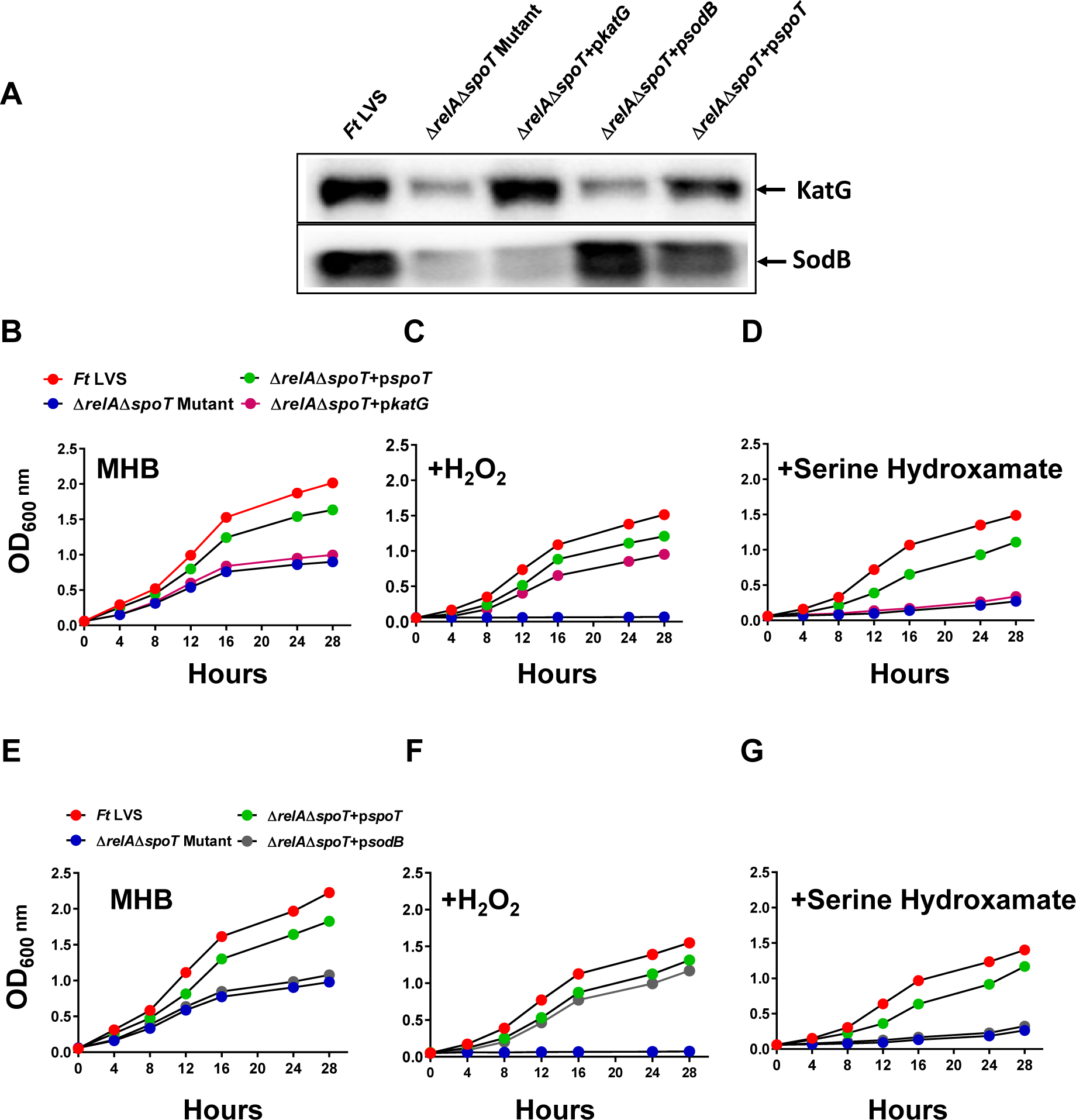
The sensitivity of the ΔrelAΔspoT mutant of F. tularensis overexpressing KatG and SodB to oxidants. (A) The western blots of the lysates of the indicated *Francisella* strains probed with anti-KatG antibodies, that were stripped and re-probed with antibodies against SodB. *F. tularensis* (*Ft*) LVS, *the* Δ*relA and* Δ*relA*Δ*spoT mutants* and the transcomplemented strains were grown in (B and E) MH-broth, (C and F) MH-broth containing 1mM H_2_O_2_ and (D and G) 500µM serine hydroxamate. The cultures were grown for 28 hours, and OD_600_ readings were recorded every 4 hours. The data are representative of two independent experiments conducted with identical results.

**Figure 9.**
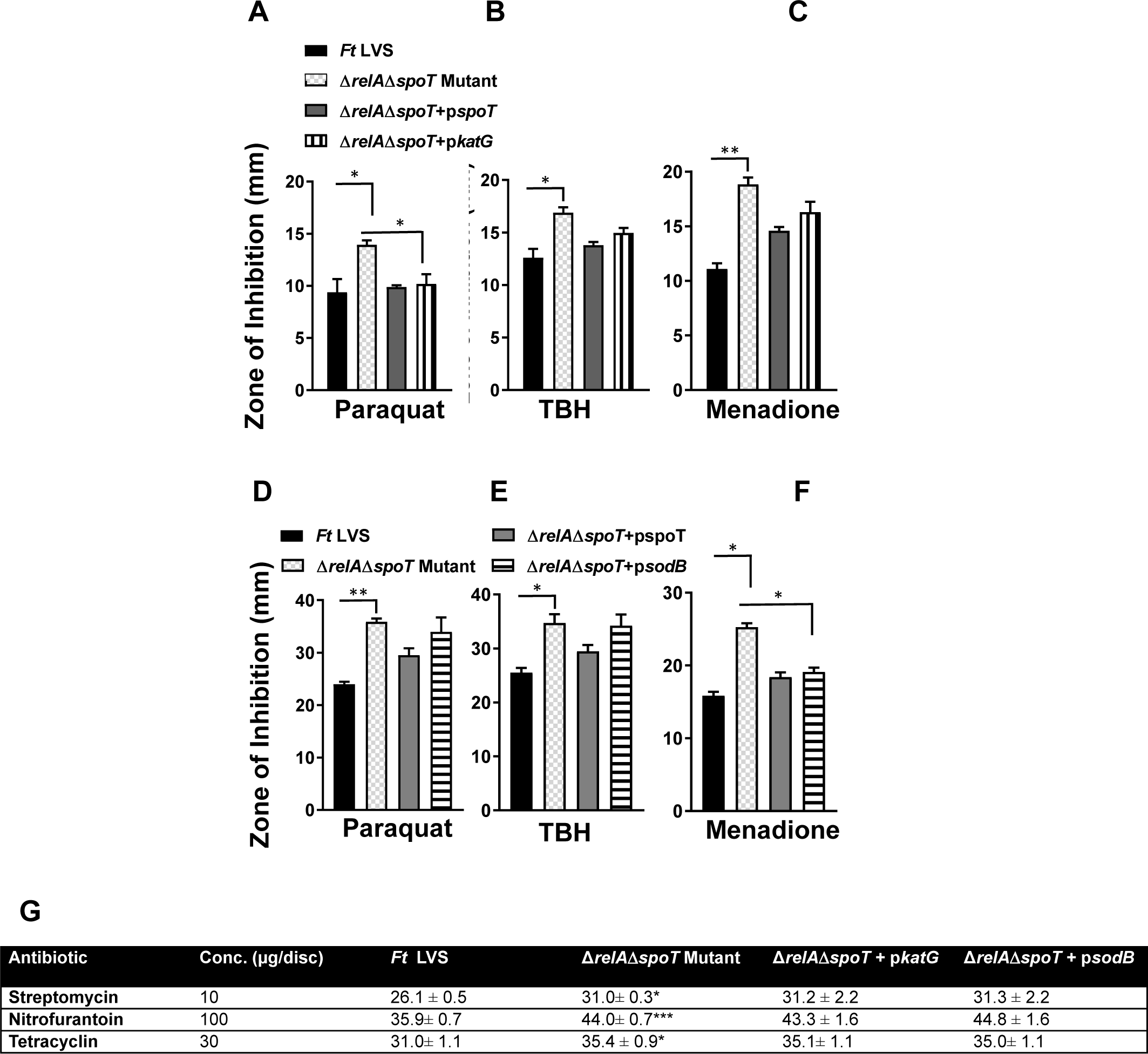
The sensitivity of the ΔrelAΔspoT mutant of F. tularensis overexpressing KatG and SodB to oxidants. Disc diffusion assay with (A and D) paraquat, (B and E) TBH and (C and F) menadione and (G) antibiotics using the indicated strains of *F. tularensis*. The data are representative of two-three independent experiments conducted with identical results. The data were analyzed by ANOVA, and the *P* values were recorded. \**P*<0.05; \*\**P*<0.01; *** *P*<0.001. Comparisons are shown with *Ft* LVS or the Δ*relA*Δ*spoT* mutant transcomplemented with *KatG* and *sodB*.

### Replication of Δ*relA*Δ*spoT* mutant of *F. tularensis* LVS is partially restored in *phox^-/-^* BMDMs

Our preceding results demonstrated that the Δ*relA*Δ*spoT* mutant is attenuated for intramacrophage growth, sensitive to oxidants and overexpression of antioxidant enzymes KatG and SodB reverts the oxidant sensitivity. We next investigated is the replication deficient phenotype of the Δ*relA*Δ*spoT* mutant is restored by infecting *phox^-/-^* macrophages which are incapable of generating reactive oxygen species. It was observed that nearly 15-fold higher Δ*relA*Δ*spoT* mutant bacteria were recovered from the *phox^-/-^*macrophages than those recovered from the wild type macrophages, indicating that the mutant bacteria can replicate in the absence of oxidative stress (Fig. 10). These results demonstrate that the stringent response mediated by RelA/SpoT of *F. tularensis* is required to overcome the oxidative stress in macrophages to establish its intracellular replicative niche.

**Figure 10:**
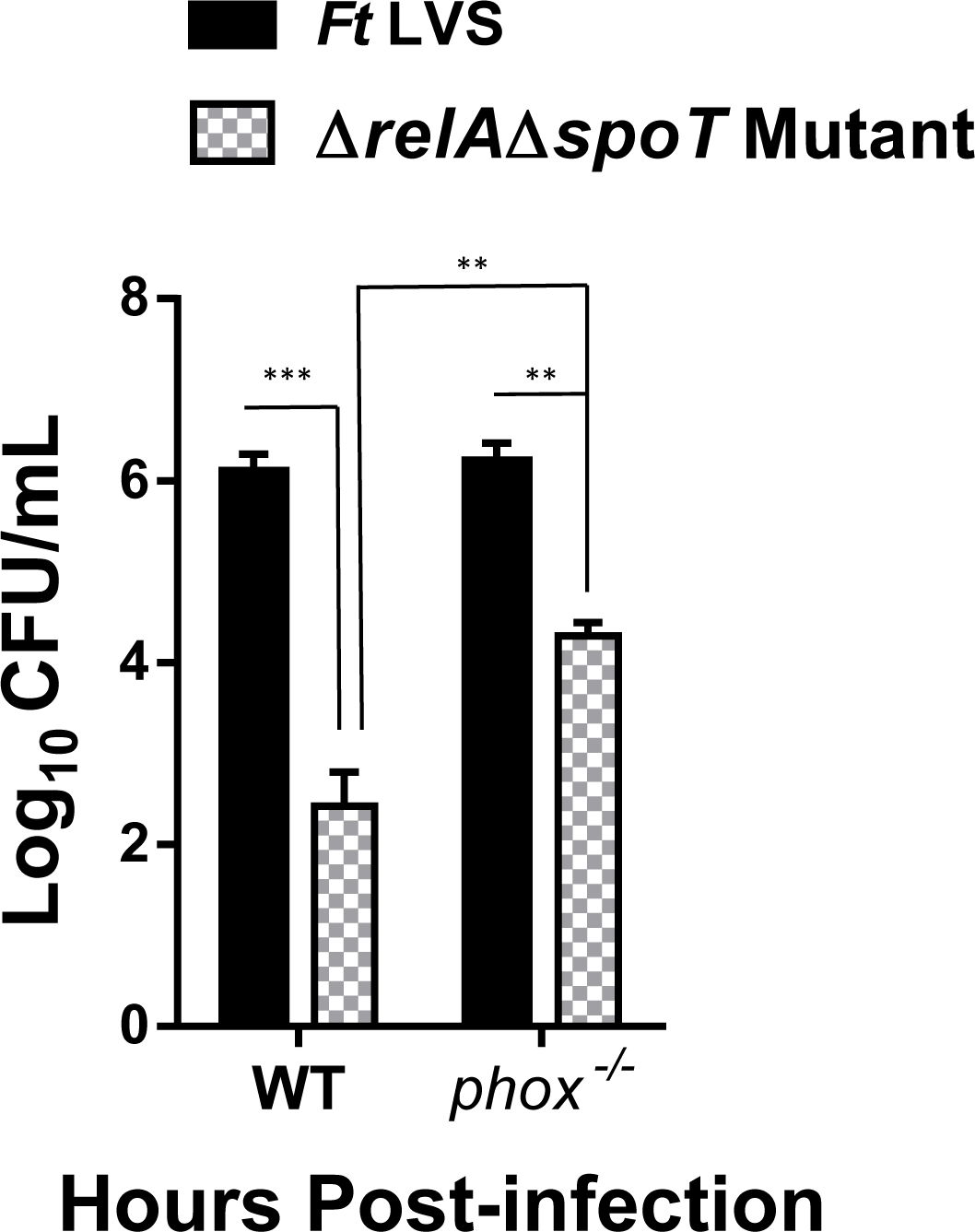
Replication of the ΔrelAΔspoT mutant of F. tularensis in phox^-/-^macrophages. Primary BMDMs derived from wild type C57BL/6, or *gp91phox^-/-^*mice were infected with the *F. tularensis* (*Ft*) LVS or the *ΔrelAΔspoT mutant* at 100 MOI (n=3 biological replicates). The cells were lysed at 4 and 24 hrs and plated on MH-chocolate agar plates for enumeration of bacterial CFU. The data are representative of three independent experiments and were analyzed using Student t-test, and *P* values were determined. \*\**P*<0.01; \*\*\**P*<0.001.

## Discussion

The synthesis of ppGpp by RelA in Gram-negative bacteria on the stalled ribosome diminishes the synthesis of stable RNA and ribosomal proteins, reprogram transcription machinery by inducing the expression of alternative sigma factors and enhance their association with core RNA polymerases. This response described as stringent response alters the half-life of RNA polymerase –promoter complexes to up- or downregulate the expression of several genes (16). The SpoT is a bifunctional synthetase and hydrolase that regulates the intracellular concentrations of (p)ppGpp. The SpoT-dependent accumulation of ppGpp occurs under carbon, iron, and fatty acid starvation conditions (15). The SpoT in-turn is regulated by ribosome-associated proteases and acyl carrier proteins (24, 25). In *Francisella,* although not mechanistically demonstrated, it has been proposed that SpoT is regulated by *migR, trmE,* and *chpA* genes (19). These genes, through unidentified mechanisms are involved in SpoT-mediated ppGpp production, and regulation of FPI encoded virulence-associated genes. An additional protein mediates the promoter interaction that affects the gene transcription in *E. coli* is known as DksA (26). The DksA homolog is absent in *F. tularensis*.

A very limited number of studies conducted to date on the stringent response of *Francisella* have primarily been focused on understanding the regulation of virulence gene expression encoded on the FPI. The Δ*relA* mutant of *F. novicida* fails to produce ppGpp under the conditions of amino acid starvation but grows better than the wild type strain at different stages of growth, including the stationary phase (17). Moreover, the Δ*relA* mutant of *F. novicida* is required for resistance against heat stress and virulence in mice. The expression of virulence genes requires the association of MglA and SspA proteins with RNA polymerase to form a complex which acts in conjunction with a DNA binding protein known as PigR. Additionally,

RelA/SpoT-mediated production of ppGpp is required for stable interactions of PigR with MglA-SspA-RNA polymerase complex for the expression of virulence genes (18). Similarly, MigR, TrmE, and CphA proteins of *F. tularensis* are required for accumulation ppGpp and virulence gene expression (19). Transcriptome analysis of *F. tularensis* SchuS4 under starvation-like conditions induced artificially by exposing *Francisella* to serine hydroxamate has shown upregulation of FPI encoded genes involved in virulence, stress response, and metabolism and downregulation of genes involved in transport and cell division (20). Collectively, this handful of studies have demonstrated the association of nutritional stress with virulence gene expression in *Francisella*.

Herein, we report a critical role of the stringent response in oxidative stress resistance of *F. tularensis*. The unique intracellular lifestyle of *Francisella* in addition to the nutritional stress, also exposes bacteria to oxidative stress. However, how stringent response governs the oxidative stress response of *F. tularensis* is not known. To address this question, we adopted a genetic approach and generated a single gene deletion mutant Δ*relA* and a Δ*relA*Δ*spoT* double gene mutant. Unlike previous studies that mostly used amino acid starvation as a means to induce stringent response, in this study, we investigated the role of stringent response during the exponential stage of bacterial growth in a nutritionally rich environment and when the bacteria are exposed to oxidative stress. We observed that ppGpp production was reduced drastically in both the Δ*relA* and the Δ*relA*Δ*spoT* mutants as compared to wild type *F. tularensis* LVS even when the bacteria were still in the exponential phase of growth and not exposed to any stress, indicating that expression of ppGpp is also required under homeostatic growth conditions. Our initial characterization revealed that the Δ*relA*Δ*spoT* but not the Δ*relA* mutant has a growth defect, enter stationary phase early, and exhibit high resistance towards streptomycin, nitrofurantoin, and tetracycline. These phenotypic attributes although have been reported for the Δ*relA*Δ*spoT* mutants of several other Gram-negative bacteria but have not been reported for the Δ*relA*Δ*spoT* mutant of *F. tularensis* LVS.

Our results show that a large number of genes are regulated by RelA/SpoT-dependent ppGpp even when bacteria are growing in a rich environment and not exposed to any nutritional stress. The majority of the genes that are regulated under the homeostatic growth conditions included those involved in metabolism, transport, and FPI genes. A number of genes encoding hypothetical proteins were also both up-and down-regulated in the Δ*relA*Δ*spoT* mutant. These results indicate that stringent response plays an essential role in the growth and survival of *F. tularensis* even in a nutritionally rich environment. When the bacteria were exposed to oxidative stress, twice the number of genes belonging to similar categories were differentially expressed in the Δ*relA*Δ*spoT* mutant indicating that stringent response regulates the oxidative stress response experienced by the bacteria in a nutritionally rich environment.

*Francisella* encounters oxidative stress in phagosomes as well as in the cytosol of the phagocytic cells during its intracellular residence. The FPI genes *iglA, iglB, iglC,* and *iglD* were all downregulated in the Δ*relA*Δ*spoT* mutant as compared to the wild type *F. tularensis* LVS under normal growth conditions and their expression levels did not change further when the bacteria were exposed to oxidative stress. A further downregulation of transcript levels of *pdpE, pdpC, iglJ, iglI, dotU, iglH, iglG* and *iglF, vgrG, iglE, pdpB, pdpA* encoded on two separate FPI operons under oxidative stress conditions from those observed in the untreated Δ*relA*Δ*spoT* mutant indicate that stringent response does play a role in regulation of these FPI genes when bacteria are exposed to oxidants. These results suggest that expression of *iglA, iglB, iglC,* and *iglD* that form the contractile sheath tube of the T6SS which is under the control of the stringent response, does not change upon exposure to the oxidative stress. However, the expression of genes that constitute the membrane complex of T6SS such as *pdpB, iglE* and *dotU* and the genes that encode T6SS effector proteins such as *pdpA, vgrG, iglG,* and *iglF* are upregulated when the bacteria sense oxidative stress conditions. These results indicate that under oxidative stress conditions, the RelA/SpoT-dependent ppGpp selectively upregulate FPI genes essential for the disintegration of the phagosomal wall and phagosomal escape to ensure that bacteria get released rapidly from the phagosomes into the cytosol to initiate its replication.

It was observed that expression of the FPI genes is significantly downregulated in the Δ*relA*Δ*spoT* mutant than that observed for the Δ*relA* mutant. Coherent with the expression profile of the FPI genes, the intramacrophage growth of the Δ*relA* mutant was not affected. However, the Δ*relA*Δ*spoT* mutant was severely attenuated for intramacrophage growth. These results demonstrate the contribution of SpoT-dependent production of ppGpp in FPI gene regulation and intramacrophage survival. However, both the Δ*relA* and the Δ*relA*Δ*spoT* mutants were highly attenuated for virulence upon intranasal infection in C57BL/6 mice. A previous study has reported the attenuation of Δ*relA*Δ*spoT* mutant of *F. tularensis* LVS in BALB/c mice upon intradermal challenge (18).

Similar to the FPI genes, the general stress proteins were also differentially expressed in the Δ*relA*Δ*spoT* mutant of *F. tularensis* LVS and three transcriptional profiles emerged when the bacteria were exposed to oxidative stress. The *clpB, hsp90,* and *hsp40* genes involved in disaggregation and reactivation of the aggregated proteins were downregulated in the Δ*relA*Δ*spoT* mutant only under the conditions of the oxidative stress. On the other hand, the other stress response genes that serve as disaggregation or degradation machines and are transcribed as operons (*clpP, clpX,* and *lon*; *hslU, hslV; grpE, dnaK,* and *dnaJ*) were downregulated under the normal growth conditions and got downregulated further in Δ*relA*Δ*spoT* mutant when bacteria were exposed to oxidants. The differential expression of these stress response genes was also reflected in the enhanced sensitivity of the Δ*relA*Δ*spoT* mutant towards high temperature. However, contrary to the Δ*relA* mutant of *F. novicida* (17), the viability of the Δ*relA* mutant of *F. tularensis* LVS is not affected by exposure to a higher temperature and remains similar to its wild type counterpart. The stress-related genes *usp, sohB and starvation protein A* involved in response to DNA damage remained downregulated Δ*relA*Δ*spoT* mutant in normal as well as oxidative stress conditions. These results indicate that RelA/SpoT-dependent stringent response regulates the expression of the stress response genes in the presence of oxidative stress to facilitate bacterial survival.

Several MglA-regulated genes were downregulated in the Δ*relA*Δ*spoT* mutant and remained similarly downregulated under oxidative stress conditions indicating that positive regulation of the MglA-dependent genes by RelA/SpoT is independent of the oxidative stress. A number of transcriptional regulators are also regulated both positively and negatively by RelA/SpoT. The positive regulation of the *fevR, cphA, trmE,* and *sspA* genes is independent of the oxidative stress. On the other hand, a *LySR* family transcriptional regulator (*FTL_1193*), *fur* and *oxyR* are regulated positively by RelA/SpoT only under oxidative stress conditions. However, the transcription of *migR* and *mglA* genes is independent of RelA/SpoT. The expression of several transcriptional regulators, including *pmrA,* is negatively regulated by RelA/SpoT, and this regulation is independent of the oxidative stress response. An upregulation of *pmrA1* has also been reported in bacteria grown in Brain Heart Infusion broth, which mimic intramacrophage growth conditions (27). The upregulated expression of *pmrA,* specifically in the Δ*relA*Δ*spoT* mutant is intriguing and need an additional investigation.

RelA/SpoT positively regulated a number of genes encoding antioxidant enzymes in *F. tularensis*. The most prominent ones being the *methionine sulfoxide reductase B* (*msrB*), *thioredoxin, glutaredoxin 1* and *3* under normal and oxidative stress conditions, and *katG* and *sodB* genes under normal growth conditions. However, it appears that expression of *sodC* and *ahpC* is negatively regulated by RelA/SpoT under the conditions of oxidative stress. In *E. coli,* the *trx1* and *grx2* genes are positively regulated by ppGpp during the stationary phase of growth (28). Our results demonstrate that stringent response also modulates the oxidative stress response under normal as well as oxidative stress conditions. Superoxide dismutase and catalase activities are regulated by the stringent response in *Pseudomonas aeruginosa* (29, 30). We observed that in addition to the transcript levels, the protein levels of primary antioxidant enzymes SodB and KatG of *F. tularensis* were reduced drastically in both the Δ*relA* and Δ*relA*Δ*spoT* mutants. Associated with a reduction in levels of these two antioxidant enzymes, the Δ*relA*Δ*spoT* mutant exhibited enhanced sensitivity towards peroxides and superoxide-generating compounds, which was restored by overexpressing either the SodB or KatG in the Δ*relA*Δ*spoT* mutant. These observations indicate that stringent response universally regulates oxidative stress response in Gram-negative bacteria. The regulation of the antioxidant enzymes occurs at the level of transcription controlled by ppGpp probably by regulation of oxidative stress response regulator OxyR. Expression of both *sodB* and *katG* genes is regulated by OxyR in *F. tularensis*. We observed that overexpression of either KatG or SodB only restored the wild type phenotype of the Δ*relA*Δ*spoT* mutant against some (paraquat and TBH with KatG, and menadione with SodB), but not all the oxidants tested. This implies that both these enzymes are required together to overcome the oxidative stress in the Δ*relA*Δ*spoT* mutant. Only a partial restoration of the growth of Δ*relA*Δ*spoT* mutant in the *phox^-/-^* macrophages that cannot produce the ROS also support these observations. Overexpression of both the enzymes in the Δ*relA*Δ*spoT* mutant was not performed in this study. Overexpression of the antioxidant enzymes, however, did not restore the growth of the Δ*relA*Δ*spoT* mutant under starvation-like conditions induced by growth in the presence of serine hydroxamate. These results indicate that regulation of the starvation response mediated by RelA/SpoT is independent of the oxidative stress response.

An association has been demonstrated between the SOD activity and the stringent-response-mediated multidrug resistance in *P. aeruginosa* (29, 30). The stringent response during stationary phase reduces the permeability of the cell envelop to a number of antibiotics and SOD activity contributes to this, resulting in multidrug resistance of *P. aeruginosa*. The mutants of *P. aeruginosa* in stringent response exhibit reduced SodB and catalase activities and complementation with SOD restores the antibiotic resistance. It has also been shown that the stringent response suppresses the central metabolism and facilitates the accumulation of intracellular free iron, which serves as a source of reactive oxygen species (31). However, in our study, we did not observe any change in sensitivity of the Δ*relA*Δ*spoT* mutant complemented either with KatG or SodB towards the antibiotics, and it remained similar to that observed for the Δ*relA*Δ*spoT* mutant. The reasons could have been the manner in which the experiments were conducted. In the former studies, the antibiotic sensitivity of the stringent response mutants and the transcomplemented strains were determined during the stationary phases of growth. In the current study, the antibiotic sensitivities were determined using disc diffusion assays and bacteria were grown on nutrient-rich MH-chocolate agar plates. It is probable that if the Δ*relA*Δ*spoT* mutant of *F. tularensis* transcomplemented with KatG or SodB during the stationary phase of growth may show an antibiotic susceptibility pattern similar to that observed for the wild type strain.

To conclude, the unique intracellular lifestyle of *Francisella* in addition to the nutritional stress also exposes bacteria to oxidative stress. This study demonstrates how stringent response governs the oxidative stress response of *F. tularensis*. This study further enhances our understanding of the intracellular survival mechanisms of *F. tularensis*.

## Materials and Methods

### Bacterial strains and cultures

*F. tularensis* subspecies *holarctica* live vaccine strain (LVS; ATCC 29684; American Culture Collection, Rockville, MD) obtained from BEI Resources, Manassas, VA was used in this study. Bacterial cultures were grown on modified MH-chocolate agar plates (MMH) supplemented with 2% IsoVitaleX. For broth cultures, bacteria were grown in Muller-Hinton Broth (MHB). Working stocks of all bacterial cultures were prepared by growing to mid-log phase at 37°C with 5% CO_2_ in MHB, snap-frozen in liquid nitrogen, and stored at −80°C until further use. *Escherichia coli* DH5-α strain (Invitrogen) used for cloning was grown either on Luria-Bertani (LB) broth or LB agar plates (Invitrogen). For the selection of transformants, mutants or transcomplemented strains, LB-broth or LB-agar media was supplemented with kanamycin (10mg/mL) or hygromycin (200mg/mL). The bacterial strains used in this study are shown in Table 1.

**Table 1:**
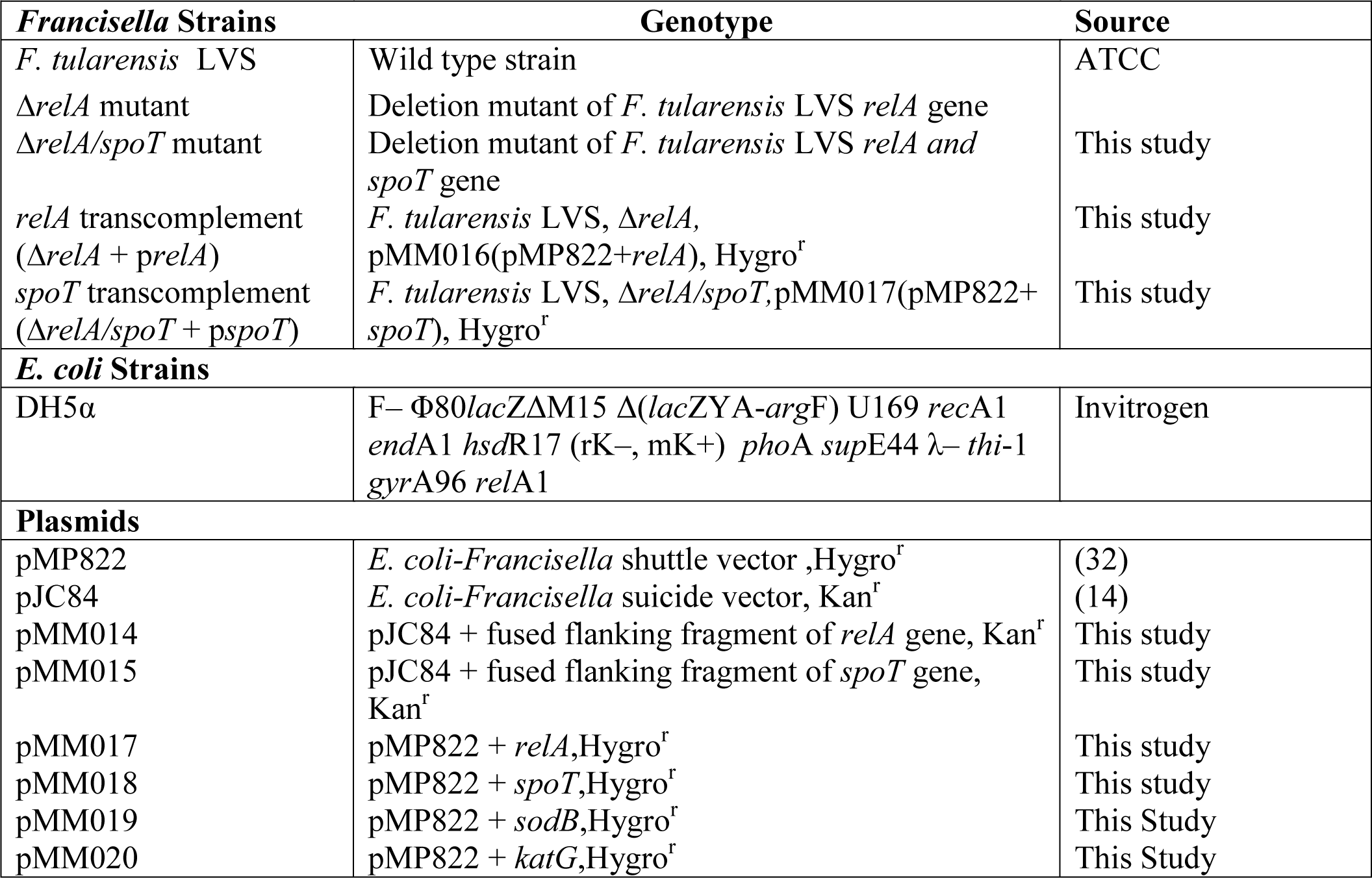
List of bacterial strains and plasmid vectors used in this study.

### Generation of *F. tularensis* Δ*relA* and Δ*relA*Δ*spoT* gene deletion mutants and transcomplemented strains of *F. tularensis* LVS

In frame single gene deletion mutants of the *relA* gene (*FTL_0285*) (Δ*relA*) and a double gene deletion mutant of *relA* and *spoT* gene (*FTL_1413*) (Δ*relA*Δ*spoT*) of *F. tularensis* were generated using a previously described allelic replacement method (10, 21, 22). For confirmation of the mutants, a duplex colony PCR was performed using internal *relA* and/or *spoT* gene-specific primers along with *sodB* gene-specific primers as internal controls. *In frame* deletions of both the *relA* and *spoT* genes was confirmed by DNA sequencing of the flanking regions. The deletion strain was designed in a way to preserve the downstream open reading frames to avoid any potential polar effects due to gene deletions.

The Δ*relA* gene deletion mutant was transcomplemented with full-length *relA* gene cloned in the *BamHI* and *Xhol* restriction sites of *E. coli-Francisella* vector pMP882. The Δ*relA*Δ*spoT* was transcomplemented with a full-length *spoT* gene cloned in the pMP822 vector in a similar fashion. The transcomplementation vectors were electroporated into the *F. tularensis relA* and Δ*relA*Δ*spoT* gene deletion mutants and selected on MMH agar supplemented with 200µg/mL hygromycin. The transcomplementation was confirmed by PCR. The primer sequences used for generation and screening of mutants and transcomplementation are shown in Table 1.

### Detection of ppGpp by high-performance liquid chromatography (HPLC)

The ppGpp extraction was performed as previously described (23). Briefly, the bacterial cultures of the wild Type *F. tularensis* LVS, the Δ*relA* and the Δ*relA*Δ*spoT* mutant strains were grown to an OD_600_ of 0.5 at 37^ο^C in 10 ml MHB. Five ml of cultures were directly mixed with 0.5 ml of 11M formic acid followed by freezing in liquid nitrogen. The mixtures were thawed and kept on ice for 30 minutes. One ml aliquots were centrifuged at 4^ο^C at 14000 RPM for 5 minutes. The supernatants were then filtered through 0.2µM filters and stored at −80^ο^C for HPLC analysis.

The ppGpp levels were quantified by HPLC. Isocratic XBridge BEH HILIC C18-HPLC column (5μm, 4.6×150 mm) was used to determine the retention time and absorbance spectrum of a ppGpp standard (TriLink Biosciences). The presence of degradation products ppGpp or (p) ppGpp were apparent at ∼2.01 min retention time. The column was run with the buffer containing 0.36 M NH_4_H_2_PO_4_ pH 3.4, 2.5% acetonitrile at 26 °C with a flow rate of 1.5 mL/min for 30 mins. Ten μl of the supernatant was injected, and ppGpp was identified as a peak which eluted at ∼2.01 min.

### Bacterial growth curves

Wild type *F. tularensis* LVS, the Δ*relA* mutant, the Δ*relA*Δ*spoT* mutant and the corresponding transcomplemented strains were grown on MH chocolate agar plates. The bacterial cultures were then resuspended to an OD_600_ of 0.05 in MHB. The cultures were either left untreated or treated with 500μM serine hydroxamate, 0.5 and 1mM hydrogen peroxide (H_2_O_2_). All bacterial cultures were incubated at 37°C with shaking (150 rpm), and the OD_600_ values were recorded for 28 hours.

### Disc diffusion assays

For disc diffusion assays, bacterial cultures of the wild type *F. tularensis* LVS, the Δ*relA* mutant, the Δ*relA*Δ*spoT* mutant, and the corresponding transcomplemented strains grown on MH-chocolate agar plates adjusted to an OD_600_ of 2.0 in MHB. Two hundred microliters of the bacterial suspensions were spread onto MH-chocolate agar plates using a sterile cotton swab. Sterile filter paper discs were placed onto the agar and were impregnated with 10μL of paraquat (0.88μg/mL), pyrogallol (40μg/mL), menadione (0.39µg/µL), tert-butyl hydroperoxide (TBH; 21.9% solution), and cumene hydroperoxide (CHP; 1.25% solution). The plates were incubated for 72 hours at 37°C and 5% CO_2_. The zones of inhibition around the discs were measured to determine the sensitivity to the compounds tested.

### Bacterial killing assays

For bacterial killing assays, equal numbers of bacteria (1×10^9^ CFU/mL) were diluted in MHB and exposed to H_2_O_2_ (1.0mM), paraquat (1.0mM), pyrogallol (1.0mM), or a temperature of 37 or 48°C. Both treated and untreated bacterial suspensions were allowed to incubate for 1 hour and 3 hours. The cultures were then serially diluted 10-fold in PBS and plated on MH-chocolate agar plates and incubated for 72 hours at 37°C with 5% CO_2_. Viable bacteria were enumerated by counting the colonies and were expressed as Log_10_ CFU/mL.

### Macrophage cell culture assays

RAW macrophage cell line and Bone Marrow-Derived Macrophages (BMDMs) isolated from wild type C57BL/6 mice were infected with wild type *F. tularensis* LVS, the Δ*relA* mutant, the Δ*relA*Δ*spoT* mutant, and the corresponding transcomplemented strains at a multiplicity of infection (MOI) of 100. The extracellular bacteria were killed by treating with gentamycin after 2 hours of infection to allow the intracellular bacteria to replicate for 4 or 24 hours. The cells were lysed at both 4- and 24-hours post-infection, diluted 10-fold and plated for enumeration of bacterial colonies.

### Mouse survival studies

Mice experiments were conducted at the Animal Resource Facility at New York Medical College according to approved IACUC protocols. All *in vivo* survival studies were conducted in wild type C57BL/6 mice aged 6 to 8 weeks. Before inoculation, mice were anesthetized with a cocktail of Ketamine and Xylazine. The mice were then intranasally inoculated with doses ranging from 1×10^5^ to 1×10^6^ of the wild type *F. tularensis* LVS, the Δ*relA* mutant, the Δ*relA*Δ*spoT* mutant in 20µL PBS. Mice were monitored up to 21 days post-infection for morbidity and mortality. Mice were weighed daily to monitor the infection. The survival results were expressed as Kaplan-Meier survival curves, and the data were analyzed using the Log-rank test.

### RNA sequencing

Overnight cultures of wild type *F. tularensis* LVS and the *ΔrelA*Δ*spoT* mutant were adjusted to an OD_600_ of 0.2 and were grown for 3 hours at 37^ο^C with shaking in 10 ml MH-broth in the absence or presence 1mM H_2_O_2_. The bacterial cells were pelleted, and total RNA was purified using Purelink RNA Mini Kit (Ambion). The contaminating DNA from RNA preparations was removed using TURBO DNA-free kits (Invitrogen). The RNA samples were submitted to the Genomics Core Facility at New York Medical College for RNA sequencing.

### Transcriptional analysis of the target genes

The RNA from wild type *F. tularensis* LVS, the *ΔrelA, ΔrelA*Δ*spoT* mutants and the corresponding complemented strains were isolated as described above. cDNA was synthesized using the iScript cDNA Synthesis Kit (Ambion). Quantitative real-time PCR (qPCR) was performed using iQ SYBR Green Supermix (BioRad) to determine the transcriptional levels of genes of interest. Expression of *tul4* gene was used as an internal control. Relative levels are represented as fold change and were calculated as follows: 2^−ΔΔ*CT*^ = 2^-^[^Δ*CT*^ ^(mutant)−Δ*CT*^ ^(WT)^], where Δ*C_T_* = *C_T_*target gene − *C_T_* internal control. The primer sequences used for qRT-PCR are shown in Table 2.

**Table 2.**
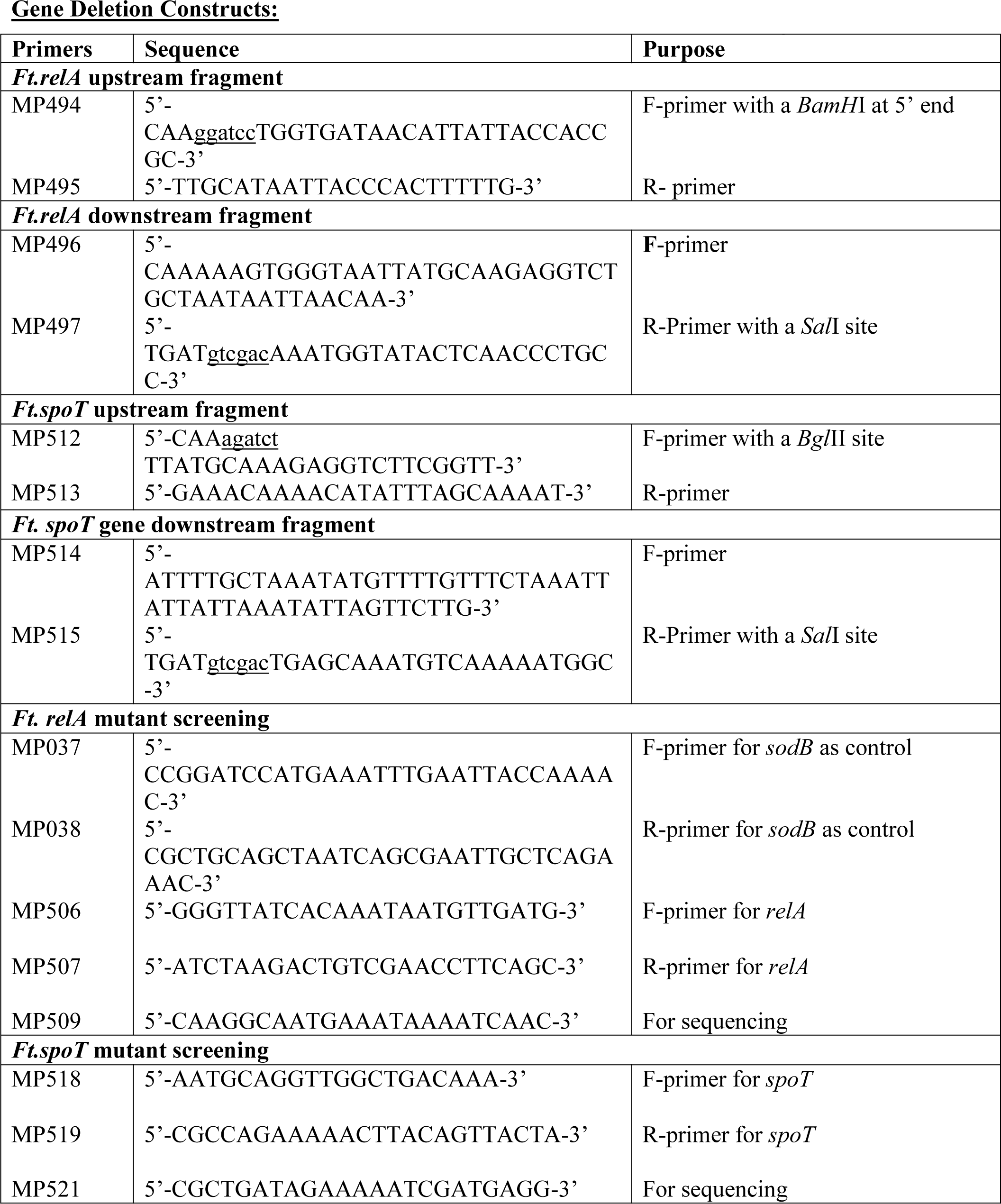
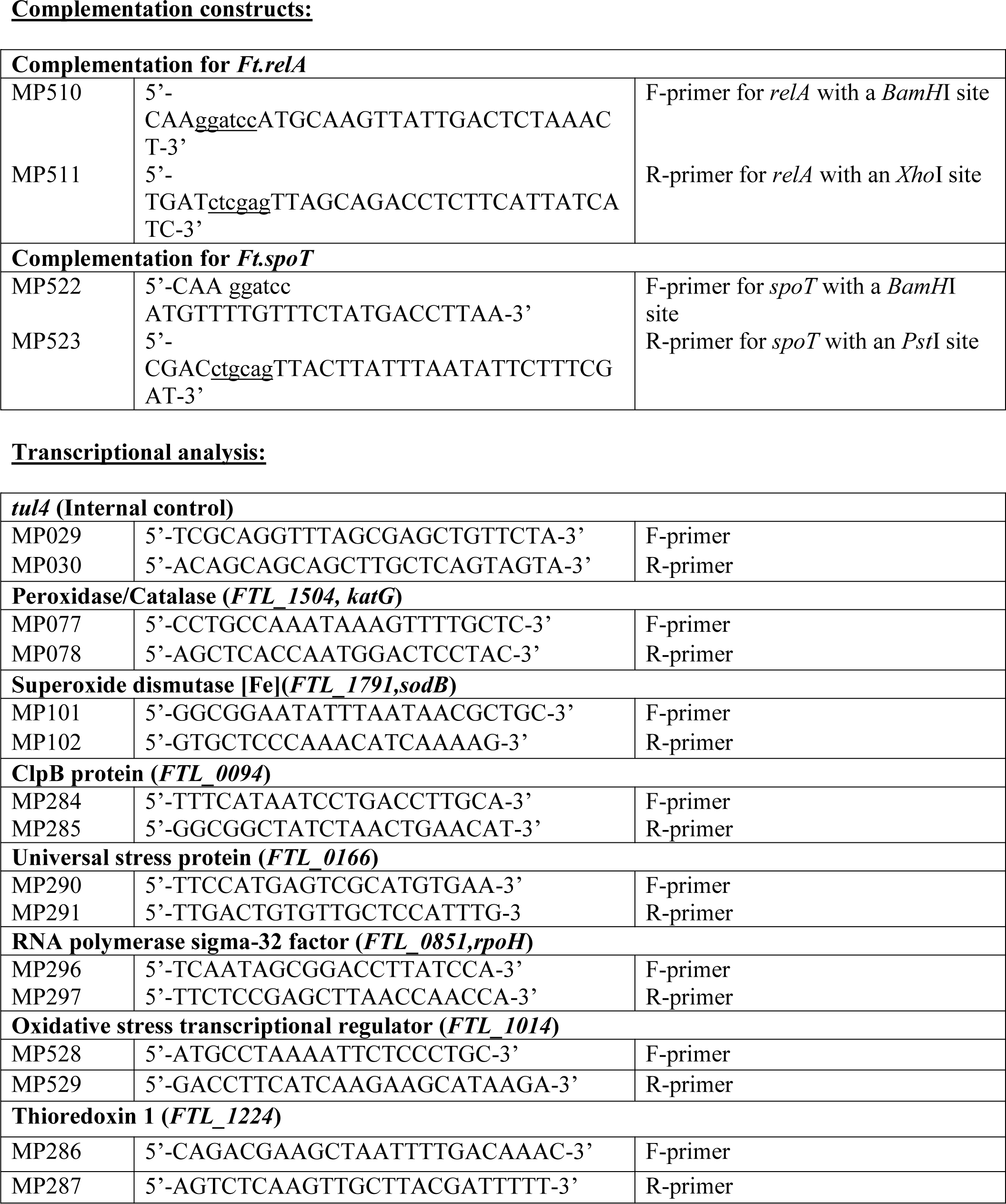
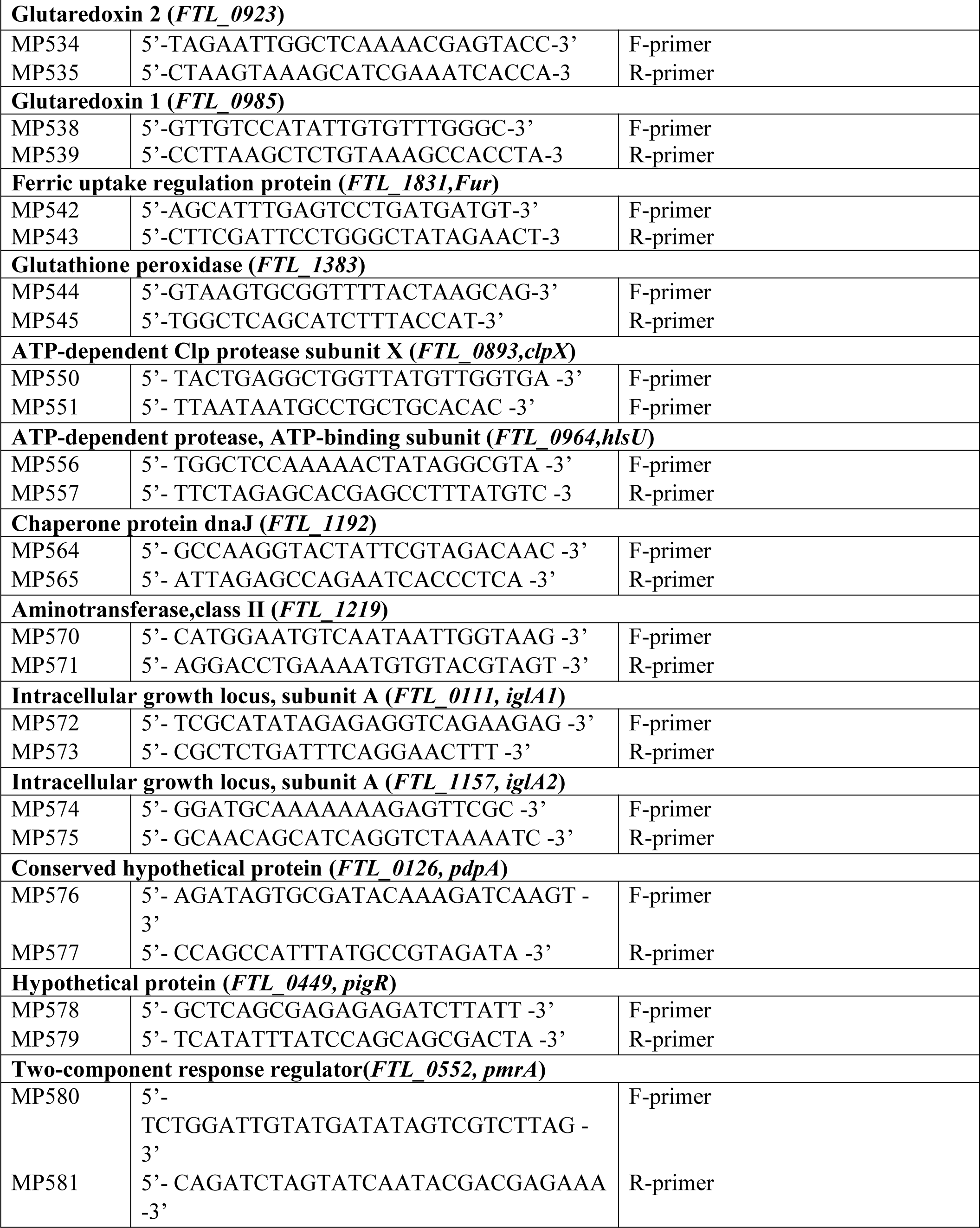
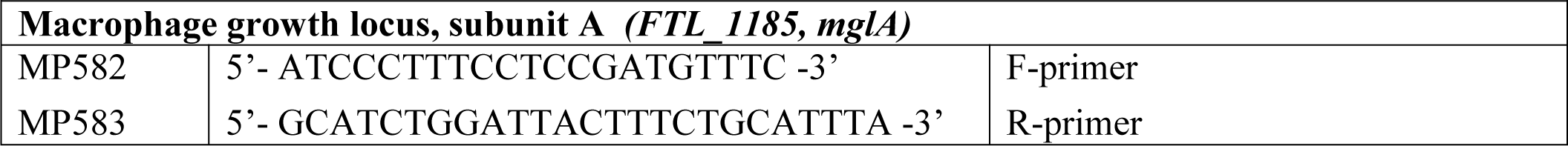
**List of primers used for gene deletion, transcomplementation and qRT-PCR**

### Overexpression of superoxide dismutase (SodB) and catalase (KatG)

To express SodB and KatG in the *ΔrelA*Δ*spoT* mutant, the full-length *sodB,* and *katG* gene sequences were amplified and cloned into pMP822 vector. The resulting plasmids were electroporated into the *ΔrelA*Δ*spoT* mutant and selected on MMH agar supplemented with 200µg/ml hygromycin. The expression of SodB and KatG proteins was confirmed by western blot analysis using anti-KatG (1:20000) and anti-SodB (1:20000) antibodies (kindly provided by Dr. Karsten Hazlett, Albany Medical College, Albany NY) and secondary monoclonal antibodies (anti-rabbit immunoglobulin, IgG,1:5000) conjugated to horseradish peroxidase (Amersham), respectively. The protein bands on the membrane were visualized using Supersignal West Pico chemiluminescent substrate (Thermo scientific) on a Chemidoc XRS system (BioRad).

### Statistical analysis

All results were expressed as means ± S.E.M. or S.D. One-Way ANOVA followed by Tukey-Kramer Multiple Comparisons tests and Student’s t-test were used for statistical analysis of the data. Survival data were analyzed using the Log-rank test and graphed using Kaplan-Meier survival curves. A *P* value of less than 0.05 was considered significant.

## Acknowledgments

This work was supported, in whole or in part, by National Institutes of Health Grants R15AI107698 (MM) and R56AI101109 (CSB). No financial conflicts of interest exist regarding the contents of the manuscript and its authors.

